# A Unique Graph-Theoretic Truth Table for Cross-Gene Branch Identity

**DOI:** 10.64898/2026.07.22.740066

**Authors:** Jiaqi Wu

**Affiliations:** Graduate School of Integrated Sciences for Life, Hiroshima University, Japan

**Keywords:** branch identity, taxon restriction, gene-tree discordance, phylogenomics, graph quotient, split, gene-by-branch matrix, missing taxa

## Abstract

Many comparative analyses operate on rectangular matrices whose columns represent the same variables across observations. Phylogenomic measurements, however, are attached to tree branches. Converting locus-specific trees into a common locus-by-coordinate matrix is straightforward only when loci contain the same taxa and compatible topologies. In real datasets, missing taxa can delete reference branches or collapse adjacent branches into composite coordinates, while gene-tree discordance can cause a coordinate that survives taxon restriction to be absent from the empirical tree. Without a formal account of these changes, non-equivalent quantities may enter the same column and distinct causes of missingness may be conflated.

Using standard tree-restriction operations, I first construct a provenance-retaining coordinate ledger. For each locus, every reference edge is recorded as deleted, retained individually, or incorporated into a composite coordinate whose original-edge membership is preserved. Each surviving coordinate is then assigned a recovery state by asking whether its corresponding split is displayed in the normalized empirical tree. Composite member sets generated across loci define a common column set, yielding one locus-by-coordinate state matrix.

The matrix is unique because the graph reduction is unique. Retained taxa span one minimal subtree; its degree-two vertices lie on uniquely determined nonbranching paths, so suppressing them in any order yields the same reduced tree and grouping of original edges. Each edge therefore has one fate, each composite coordinate one member set, and each empirical split query one answer. Thus every locus-by-coordinate cell has one determined state under fixed labelled inputs and conventions. Fiber–split equivalence shows that projected splits faithfully encode these graph-derived edge groups, explaining why split-based implementations recover the same coordinate structure and cell states.

A deterministic exhaustive validation on the six-taxon worked-example domain evaluated 14,160 primitive and composite cells across 472 normalized empirical-tree cases. Separately implemented graph- and split-side evaluators agreed in every tested cell, and two clean runs produced byte-identical canonical result files.

No branch lengths are required. When valid lengths are supplied, numerical entries may be added separately by single-edge lookup or a declared selected-component-edge sum. The theorem guarantees a unique coordinate-and-state matrix under the stated inputs, not an input-independent numerical matrix or historical truth. It provides an auditable foundation for branch-wise comparative analyses under heterogeneous taxon coverage and gene-tree discordance.

## 1. Introduction

Many modern comparative analyses begin with a rectangular matrix: observations occupy rows, variables occupy columns, and every column must retain the same meaning across observations. Phylogenomic data do not naturally arrive in this form. Evolutionary measurements such as substitution rates, selective constraints, expression changes, and other lineage-associated quantities are attached to branches of trees. Because different loci may contain different taxa or support different topologies, heterogeneous locus trees cannot automatically be treated as rows of one common gene-by-coordinate matrix. Before any joint analysis can begin, it must first be established that the quantities placed in one matrix column refer to the same evolutionary coordinate.

Consider a study that seeks to compare a branch-associated measurement across many loci. At one locus, the relevant reference branch may remain independently represented. At another, missing taxa may remove a nearby branching point and force that branch to merge with adjacent reference edges. At a third, the reference coordinate may survive taxon restriction, yet the corresponding branch relationship may be absent from the estimated gene-tree topology. Values from these situations cannot be treated as interchangeable merely because they are assigned the same nominal branch label. Nor should their empty cells be interpreted identically: structural deletion, fusion into a larger coordinate, and topological non-recovery arise from different mechanisms and carry different information about what the locus can represent.

Figure 1 illustrates this representational problem. Restricting a reference tree to the taxa retained at one locus can delete an original edge, preserve it as an individual coordinate, or collapse several original edges into one composite coordinate. The empirical gene tree enters only after this reference-level reduction. It determines whether a surviving coordinate is recovered in the empirical topology, but it cannot alter a deletion or fusion state already fixed by reference restriction. A valid locus-by-coordinate matrix must therefore preserve two kinds of information: the fate of every original reference edge under taxon restriction, and the empirical recovery state of every coordinate that remains representable.

**Figure 1.**
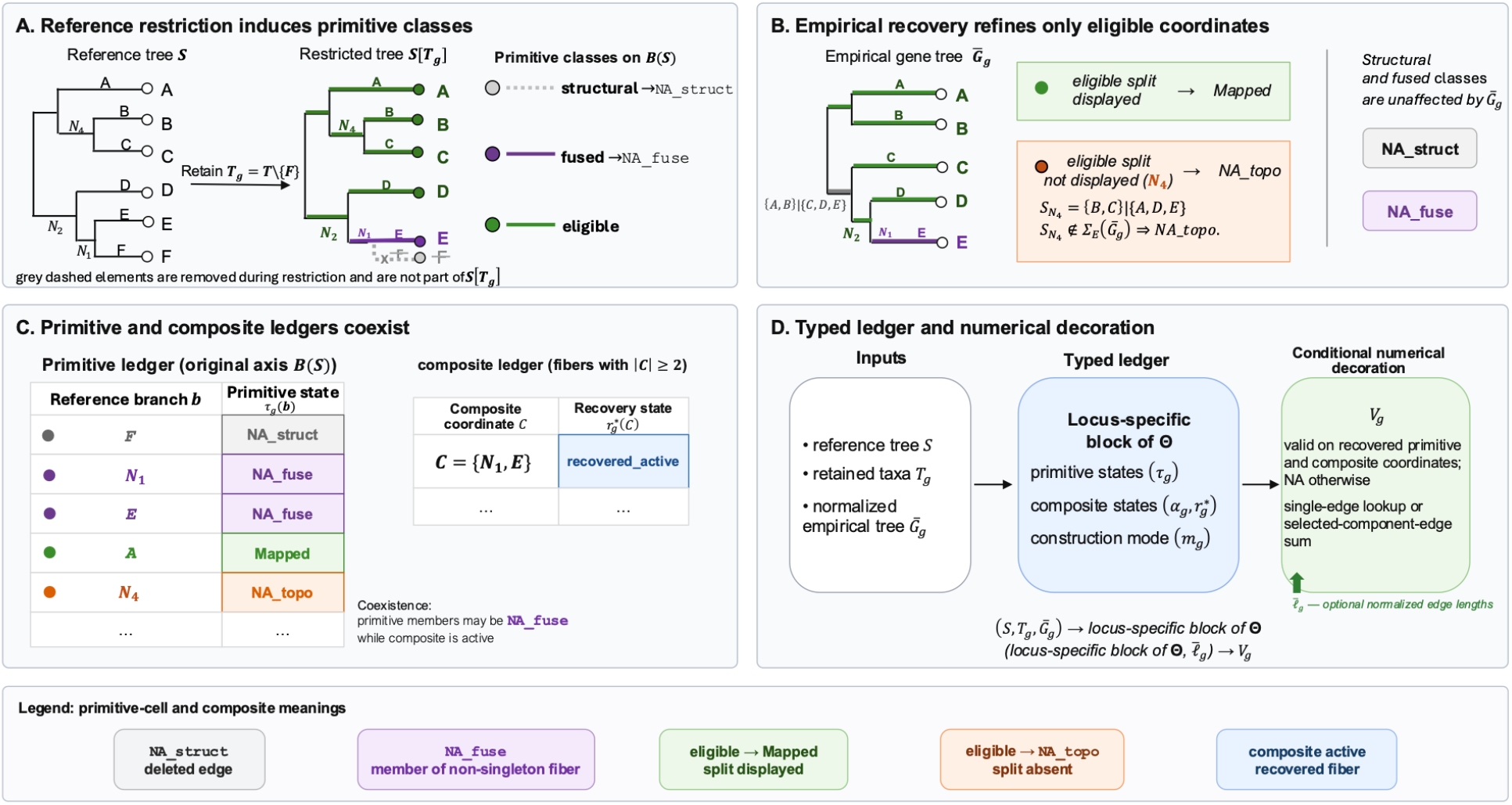
Overview of the framework: from reference restriction to a typed branch-coordinate ledger. **(A)** Restricting the labelled reference tree (*S*) to the retained taxon set (*T*_*g*_ = *T* ∖ {*F*}) deletes the terminal reference edge (*F*), which is classified as structural and assigned `NA_struct`. Suppression of the degree-two vertex exposed by this deletion contracts the internal edge (*N*_1_) and terminal edge (*E*) into the non-singleton fiber (*C* = {*N*_1_, *E*}); the primitive members are therefore assigned `NA_fuse`, whereas the remaining singleton fibers are eligible. Grey dashed elements are removed during restriction and are not part of (*S*[*T*_*g*_]). **(B)** Empirical recovery is applied only to eligible coordinates. In the normalized empirical tree 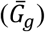, the terminal coordinate (*A*) is displayed and is therefore `Mapped`, whereas the eligible internal coordinate (*N*_4_), with restricted split ({*B, C*} ∣ {*A, D, E*}), is not displayed and is assigned `NA_topo`. Structural and fused states fixed by reference restriction are unaffected by the empirical topology. **(C)** The primitive and composite ledgers coexist. On the frozen primitive axis (*B*(*S*)), (*F*) is `NA_struct`, (*N*_”_) and (*E*) are `NA_fuse`, (*A*) is `Mapped`, and (*N*_4_) is `NA_topo`. At the same locus, the composite coordinate (*C* = {*N*_1_, *E*}) remains active and is recovered as `recovered_active`. **(D)** The fixed reference tree, retained taxa, and normalized empirical topology determine one locus-specific block of the categorical ledger (*Θ*), including primitive states, composite states, and the value-construction mode. Optional normalized edge lengths 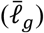 enter only afterwards to produce the conditional numerical decoration (*V*_*g*_) by single-edge lookup or a declared selected-component-edge sum.

The graph operations underlying tree restriction are classical. A phylogenetic tree restricted to a retained leaf set is obtained from the unique minimal subtree connecting those leaves, followed by suppression of degree-two vertices; edge-induced splits provide a standard representation of the resulting tree (Buneman 1971; Semple and Steel 2003; Bryant et al. 2004). Recent work has also extended split-system representations to partial splits and subtree distances (Bryant et al. 2025). These classical constructions determine restricted trees and their displayed splits, but they do not retain the frozen-edge provenance needed to assemble a cross-locus matrix containing both original and composite coordinates and one state for every locus-by-coordinate cell. In particular, a partial split is a bipartition of a subset of taxa, whereas the composite coordinates considered here are exact sets of original reference edges generated by restriction and degree-two suppression.

Here, I construct this provenance-retaining coordinate ledger and prove its fixed-input determinacy. For each locus, the retained taxa determine a unique minimal subtree of a fixed labelled reference tree. The remaining degree-two vertices lie on uniquely determined paths with no further branching, so suppressing them in any order yields the same reduced tree and the same grouping of original edges. Edges outside the connecting subtree are recorded as deleted; a singleton surviving group defines an individual coordinate; and a non-singleton group defines a composite coordinate identified by its exact original-edge membership. The finite union of the composite member sets generated across loci supplies a common cross-locus column set.

Splits enter at a separate interface. The coordinate groups are defined by graph restriction and suppression, not by a split-matching convention. Projected splits faithfully encode the already determined edge groups, and the normalized empirical gene tree is queried only to determine whether the split associated with a surviving coordinate is displayed. Empirical topology therefore refines the reference-level ledger without redefining it. This separation yields a four-state primitive output—`Mapped`, `NA_struct`, `NA_fuse`, or `NA_topo`—but these four labels are not one intrinsic graph partition. They arise from a three-state reference classification—structural, fused, or eligible—followed by a two-state empirical refinement of the eligible class.

Fusion also requires coordinates beyond the original reference-edge axis. When several original edges collapse into one surviving path, their individual rows are correctly marked `NA_fuse` because they are no longer separately identifiable on that retained-taxon set. The path itself nevertheless remains represented and must be recorded as a composite coordinate. Across loci, different restrictions may generate different, sometimes overlapping, original-edge member sets. I collect these member sets into a finite composite-coordinate ledger and determine, at each locus, whether a previously generated composite is present as one current path, exactly representable as a union of whole current paths, or unavailable because it contains a deleted edge or cuts through an indivisible current coordinate. This ledger is a catalogue of possible composite representations, not an additive or orthogonal basis.

Combining the primitive and composite assignments produces the complete locus-by-coordinate state matrix. I use the term “truth table” in an enumerative sense: it is the complete finite assignment of one graph-determined state to every cell under fixed labelled topological inputs and stated conventions. It is not a Boolean truth table and does not imply that the supplied reference tree or empirical gene trees are historically correct. The main theorem proves that, once the labelled reference tree, retained-taxon sets, normalized empirical topologies, and comparison conventions are fixed, the matrix columns and every cell state are uniquely determined.

Branch lengths are not required for this categorical result. When valid empirical edge lengths are additionally supplied, numerical entries can be attached only after the coordinate and state structure has been fixed. A recovered value is then obtained either by looking up one uniquely identified empirical edge or by applying a declared sum over selected component edges. This numerical layer is conditional on the supplied lengths and construction rule. It does not assert that an empirical fused-edge length decomposes into latent primitive contributions, that selected component edges form a continuous empirical path, or that any supplied length is an evolutionary ground-truth quantity.

The theory is related to, but distinct from, the SplitAligner software and its empirical audits (Wu 2026). SplitAligner provides an operational split-based construction of gene-by-coordinate matrices. The present study establishes the graph-theoretic object that this construction recovers and proves why its coordinate and state assignments are single-valued under fixed inputs. The agreement between graph-based restriction and split-based recovery is not an empirical coincidence between algorithms: it follows because projected split classes faithfully encode the same restriction-induced original-edge groups. Production implementation, runtime behavior, large empirical applications, and downstream consequences of invalid coordinate representations remain the subjects of the companion methodological and audit studies.

The remainder of the paper develops this result in layers. Section 2 defines the reference and empirical tree inputs and introduces the running example. Section 3 constructs the reference-level restriction ledger. Section 4 adds empirical recovery only for coordinates that remain eligible after restriction. Section 5defines the cross-locus composite-coordinate universe and its locus-specific representations. Section 6 assembles the unique coordinate-and-state matrix and its separate conditional numerical decoration. Section 7 reports an exhaustive computational validation of the translation from the mathematical definitions to executable state assignments. The Discussion considers the representational implications of the theorem and the relationship between graph-based and split-based constructions.

## 2. Formal setting

I distinguish the fixed reference coordinate system from the empirical trees used to recover it. This separation is essential: taxon restriction determines the categorical reference object, whereas empirical topology and branch lengths enter only in later layers.

### 2.1 Reference tree and locus-specific taxon sets

Let *S* be a finite, simple, connected, acyclic, unrooted, binary tree. Its degree-one vertices are bijectively labelled by the taxon set *T*, its primitive edge set is *B*(*S*), and every internal vertex has degree three. Here, *primitive* means atomic on this frozen reference edge axis. It does not imply that an edge remains independently identifiable after every taxon restriction. Nor does it assert that the same primitive axis would be retained if the reference tree or taxon universe were replaced. The binary assumption is imposed to make the chosen reference resolution coordinate-complete: every relationship represented by that resolution has an explicit primitive edge address. A multifurcating reference tree would still determine the restriction quotient and categorical states on its existing edges, but any unchosen refinement at a polytomy would be absent from *B*(*S*), not classified as `NA_topo`. Binary resolution is therefore a scope choice for the reference coordinate universe, not a claim that *S* is historically correct or a prescription to resolve an uncertain biological polytomy arbitrarily.

Let *G* be a finite locus set. Each *g* ∈ *G* has a retained taxon set

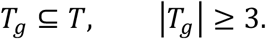

For each locus, *K*_*g*_, *D*_*g*_, *H*_*g*_, *F*_*g*_, and *π*_*g*_ denote the restriction objects defined in Section 3. The finiteness of *S* and *G* ensures that both the primitive and composite cell-index sets are finite.

### 2.2 Normalized empirical trees

An admissible raw empirical input is finite, simple, connected, and acyclic and carries the taxon labels *T*_*g*_. It may be rooted and may contain degree-two internal vertices. I forget root orientation only when the resulting unrooted graph has degree-one vertices bijectively labelled by *T*_*g*_, so an unlabelled root cannot become a leaf. Maximal chains joined through degree-two internal vertices are then suppressed to obtain 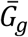.

Topological normalization does not require branch lengths. If the raw tree carries a total finite nonnegative length on every edge, each normalized edge receives the finite sum along its contracted chain. If lengths are missing, partial, or invalid, the topology remains available for categorical recovery, but the conditional numerical theorem is not invoked until a total valid normalized function 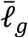 is supplied.

This normalization is unique up to leaf-label-preserving unrooted isomorphism. The resulting 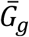 is finite, simple, connected, acyclic, has degree-one vertices bijectively labelled by *T*_*g*_, and has no degree-two internal vertex. Multifurcations are allowed. The two no-degree-two requirements have different roles: in *S*, they prevent representational subdivision of the frozen primitive reference axis; in 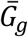, they guarantee that a displayed split has a unique empirical edge address. Suppressing a degree-two chain removes a representational subdivision, not a branching event, and preserves the labelled splits displayed by the normalized topology. By contrast with S, the normalized empirical tree is queried against an already fixed reference axis rather than used to define the coordinate universe. Empirical multifurcations are therefore permitted: they may leave an eligible reference split undisplayed, but they do not remove that reference coordinate from the ledger.

*A proof of the normalization result is given in Appendix A*.*2*.

### 2.3 Splits and canonicalization

A split is an unordered bipartition of a labelled leaf set into two nonempty blocks. For a nondeleted reference edge 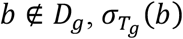 denotes the two-nonempty-block split that it induces on *T*_*g*_. For an empirical edge 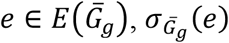 denotes its displayed split. The full empirical edge-split set is

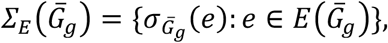

including terminal singleton splits.

All comparisons use a fixed faithful canonicalization *K*_*can*_, satisfying

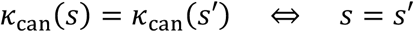

for unordered labelled splits. Canonicalization is only a collision-free notation for comparing unordered splits; it does not define branch identity. For a split set *A*, write

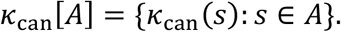

### 2.4 Optional numerical decoration

The categorical theorem requires no branch lengths. When numerical values are considered, each normalized empirical tree is additionally supplied with a total nonnegative edge-length function

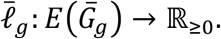

These estimated lengths decorate already determined categorical cells; they do not define branch identity or constitute historical ground truth.

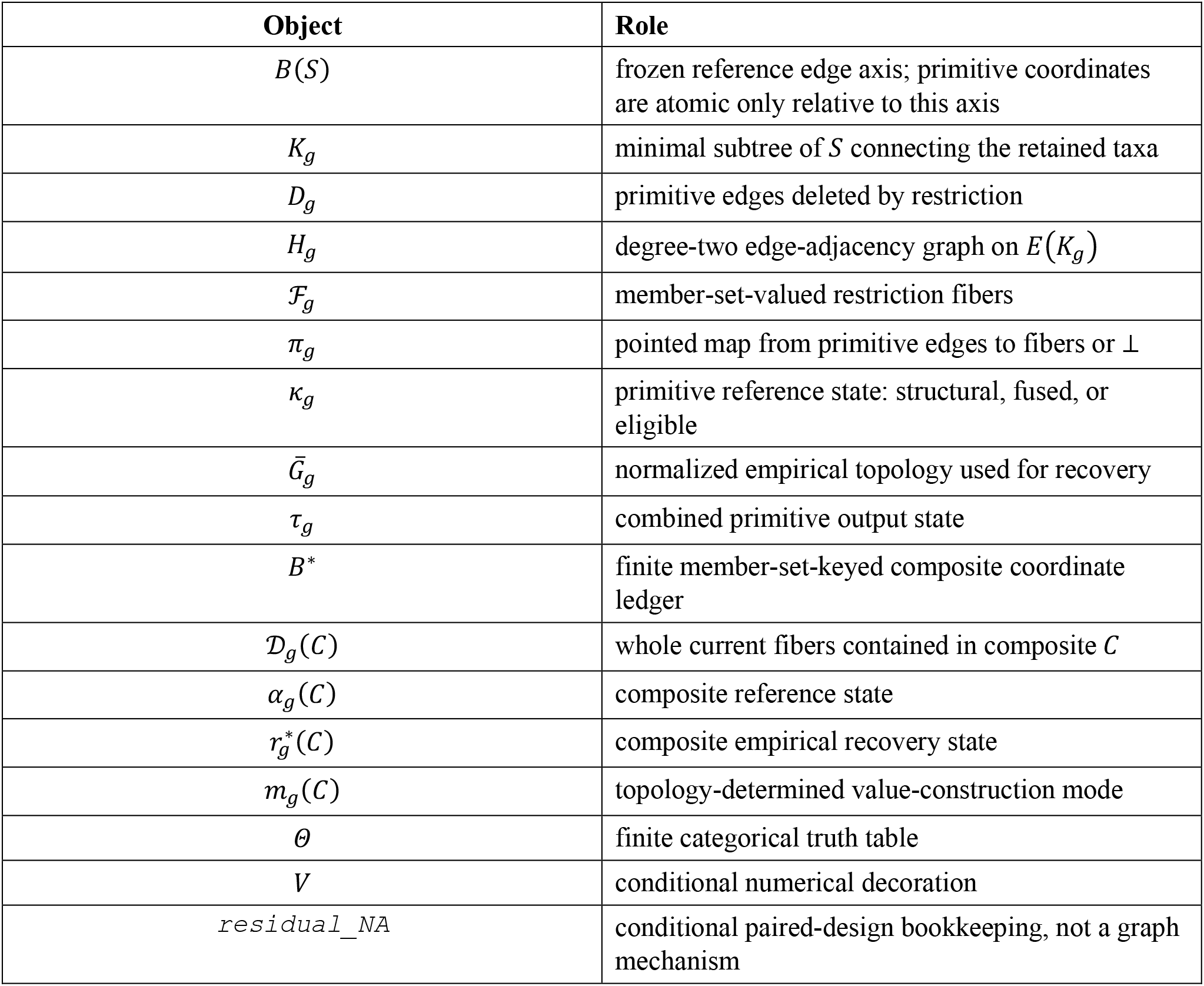

### 2.5 Running example

Figure 1 uses one six-taxon example throughout. Let *S* be the labelled unrooted reference tree displayed by the Newick string (*A*, (*B, C*), (*D*, (*E, F*)));, with taxon set *T* = {*A, B, C, D, E, F*} and reference-edge labels as shown in Figure 1A. At locus *g*, retain *T*_*g*_ = *T* ∖ {*F*} = {*A, B, C, D, E*}.

Restriction deletes the terminal reference edge *F*. Suppressing the degree-two vertex exposed by this deletion contracts the internal edge *N*_1_ and terminal edge *E* into the non-singleton fiber *C* = {*N*_1_, *E*}; all other nondeleted fibers are singletons. Thus *F* is `NA_struct`, *N*_1_ and *E* are `NA_fuse`, and *C* remains as an active composite coordinate.

In the normalized empirical topology shown in Figure 1B, the eligible terminal coordinate *A* is `Mapped`, whereas the eligible internal coordinate *N*_4_ is `NA_topo` because its split {B, C} ∣ {A, D, E} is absent. The composite *C* is `recovered_active` because the empirical tree displays the terminal split of *E*. The example therefore contains structural deletion, primitive fusion, empirical non-recovery, and recovery of the corresponding composite coordinate.

| Graph outcome | Coordinate | Reference state | Primitive or composite outcome |
| --- | --- | --- | --- |
| Edge deleted by restriction | $F$ | structural | $F$: `NA_struct` |
| Endpoint-collapse fiber | $C = \{N_1, E\}$ | $N_1, E$: fused | $N_1, E$: `NA_fuse`; $C$: `recovered_active` |
| Eligible terminal coordinate | $A$ | eligible | Mapped |
| Eligible internal coordinate | $N_4$ | eligible | `NA_topo` |

The Newick string and tree drawings are display conventions only; the labelled graphs, retained-taxon set, and original-edge membership of *C* define the example.

## 3. A canonical reference truth layer under taxon restriction

Cross-locus branch comparison starts from the fixed primitive axis *B*(*S*). Restricting taxa does more than produce a smaller tree: it determines, for every original edge, whether that coordinate is deleted, retained alone, or retained only as part of a fused path. This reference classification reads neither an empirical gene tree nor a branch length.

### 3.1 Restriction induces a pointed edge quotient

For locus *g*, let *K*_*g*_ be the minimal subtree connecting *T*_*g*_, with deleted set

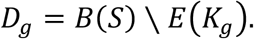

Taxon deletion may leave degree-two vertices in *K*_*g*_. Define an edge-adjacency graph *H*_*g*_: its vertices are the edges of *K*_*g*_, and two are adjacent when the corresponding tree edges meet at a degree-two vertex. The connected components form the fiber partition *F*_*g*_; each fiber is a maximal path contracted to one reduced edge.

The map

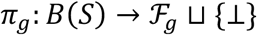

records where every original edge goes after pruning: to deletion (⊥) or to one retained fiber identified by its original-edge member set.

#### Proposition 1 (canonical pointed restriction ledger)

For fixed *S* and *T*_*g*_, *K*_*g*_, *D*_*g*_, *H*_*g*_, *F*_*g*_, and *π*_*g*_ exist and are unique, and each ordinary fiber is a connected path segment of *S*. The construction uses no gene-tree topology, branch-length estimate, or split-encoding convention; no root is involved because *S* is treated as unrooted.

*Proof: Appendix B*.*1*.

### 3.2 Projected splits encode the fibers

Lemma 1 (fiber–split interface). Every nondeleted primitive edge *b* induces an unordered split 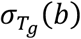. Fiber membership is equivalent to projected-split equality:

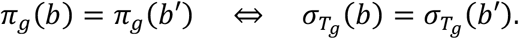

Along one degree-two fiber, moving the cut crosses no retained off-chain arm, so the split is unchanged. Conversely, if the joining chain contains a junction with a retained off-chain arm, choose retained leaves *p* and *q* beyond the two cuts and *x* in that arm; *x* is grouped with *q* under one cut and with *p* under the other, so the unordered splits differ.

Projected splits therefore encode the graph fibers but do not define them. Empirical discordance may later prevent recovery of an eligible split, but it cannot change which reference edges restriction deleted or fused.

*Proof: Appendix B*.*2*.

### 3.3 Primitive reference states

The pointed quotient gives

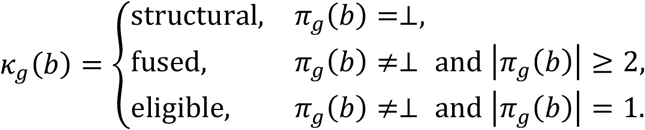

#### Proposition 2 (primitive reference trichotomy)

For every *g* ∈ *G* and *b* ∈ *B*(*S*), exactly one of structural, fused, or eligible applies. At a fixed locus, every fused primitive edge belongs to exactly one non-singleton active fiber.

*Proof: Appendix B*.*3*.

A structural edge corresponds to `NA_struct`; a fused edge corresponds to `NA_fuse`; an eligible edge is the sole member of its fiber and can proceed to empirical recovery. Fusion is not biological absence or zero length: one non-singleton fiber survives as a composite coordinate while each original member contributes a separate primitive `NA_fuse` cell.

### 3.4 Endpoint collapse

Let *b*_*x*_ be the terminal edge of a retained taxon *x*. If an internal edge *b* ≠ *b*_*x*_ also projects to

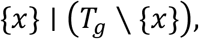

fiber–split equivalence gives *π*_*g*_(*b*) = *π*_*g*_(*b*_*x*_). Their common fiber is non-singleton, so both are fused. At the reference layer, *b*_*x*_ is eligible exactly when its fiber is {*b*_*x*_}; empirical recovery of eligible terminals is defined in Section 4.

*A proof of the endpoint-collapse result is given in Appendix B*.*4*.

### 3.5 Boundary cases

For distinct *u, v* ∈ *T*, let *C*_*uv*_ = *E*=(*P*_*s*_ (*u, v*)). Under the separate restriction to {*u, v*},

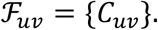

Because the frozen reference tree has at least three leaves, |*C*_*uv*_| ≥ 2. Every path member is `NA_fuse`, every off-path edge is `NA_struct`, and no primitive coordinate is eligible. This boundary construction does not alter the main, |*T*_*g*_, | ≥ 3 domain, the frozen locus family, or *B*^∗^.

*Proof: Appendix F*.*1*.

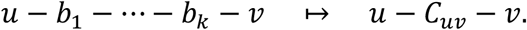

With one retained taxon, the retained-edge ledger is empty, but the fixed primitive row remains defined and uniformly `NA_struct`.

*See Appendix F*.*2*.

Together, these results establish the reference truth layer: “truth” here means one graph-determined state conditional on fixed =(*S, T*_*g*_), not historical correctness of *S* or truth of a branch-length estimate.

## 4. Empirical recovery of eligible primitive coordinates

The reference trichotomy determines which primitive coordinates can be queried in an empirical tree. Structural and fused cells are already terminal at the reference layer. Only eligible coordinates—singleton reference fibers—proceed to empirical recovery.

### 4.1 Unique empirical edge addresses

A normalized empirical tree has an injective edge-to-split map.

#### Lemma 2 (edge–split injectivity)

Let *G* be a finite, simple, connected, acyclic, unrooted tree whose degree-one vertices are bijectively taxon-labelled and which has no degree-two internal vertices. Then distinct edges of *G* display distinct unordered labelled splits.

Join two distinct edges by their unique minimal edge chain. If a junction on that chain has an off-chain component, that finite component contains a degree-one vertex and therefore, under full leaf labelling, a labelled taxon. That taxon lies on different sides of the two cuts, so the displayed splits differ. Equal splits would therefore force every junction on the joining chain to have degree two, contrary to normalization. Thus each displayed split has one empirical edge. The argument does not require the empirical tree to be binary: every internal vertex on the joining chain has degree at least three and therefore supplies an off-chain component containing a labelled leaf. This uniqueness is needed for numerical lookup; categorical presence or absence is already a Boolean split-set question.

*Proof: Appendix C*.*1*.

### 4.2 Recovery on the eligible axis

Let

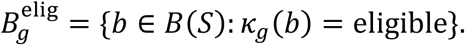

Define

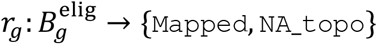

by

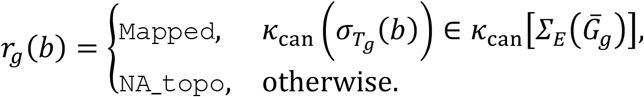

The combined primitive state is

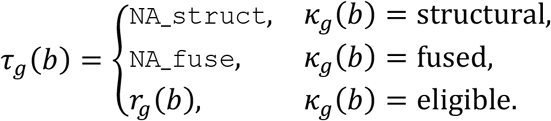

#### Proposition 3 (eligible-only empirical refinement)

For fixed = (*S, T*_*g*_, 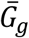) and fixed split conventions, *τ*_*g*_ is total and single-valued. Recovery refines only the eligible block; it cannot modify the deleted set, the fiber partition, or any composite coordinate defined by the reference restriction.

*Proof: Appendix C*.*2*.

This is the point at which empirical topology first enters the ledger. Here `Mapped` means only that the supplied normalized empirical topology displays the queried reference split; it validates neither tree as historical truth. A coordinate can be graph-eligible but absent from 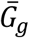, yielding `NA_topo`.

Conversely, empirical display of a split cannot turn a deleted or fused primitive coordinate into an eligible one.

### 4.3 Retained terminal coordinates

For a retained taxon *x*, the terminal reference edge *b*_*x*_ is never structural. Its complete classification is

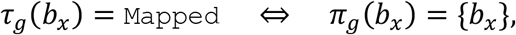

and

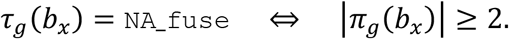

When the terminal fiber is singleton, every empirical tree on *T*_*g*_ displays the terminal split {*x*} ∣ (*T*_*g*_ ∖ {*x*}), so the coordinate is necessarily `Mapped`. When endpoint collapse occurs, the terminal edge belongs to a non-singleton fiber and is `NA_fuse` together with its internal path members. A retained terminal is therefore never `NA_struct` or `NA_topo`.

*Proof: Appendix C*.*3*.

### 4.4 Conditional paired-design bookkeeping

Some paired analyses require a deterministic residual label after the graph-derived states have been assigned. I formalize that bookkeeping separately so that it cannot be mistaken for an additional graph-theoretic state. `residual_NA` is not a fifth graph-theoretic mechanism. Let *E*_*g*_ be an explicitly stated paired-evidence domain and let

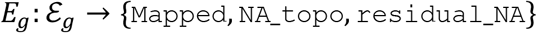

be a fixed total deterministic map. Because its inputs are already single-valued, its output is also single-valued. The conclusion is conditional on the declared *E*_*g*_. The present study proves single-valuedness for that rule but neither selects a canonical rule nor claims biological optimality. A different total deterministic evidence rule may produce a different but internally deterministic bookkeeping assignment.

*The determinacy argument is given in Appendix C*.*4*.

### 4.5 Two-taxon empirical boundary

For the separately certified two-taxon boundary coordinate *C*_*uv*_, use boundary-local maps on the singleton domain {*C*_*uv*_}:

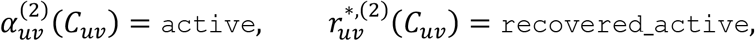

and

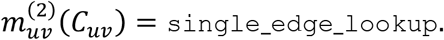

A normalized empirical tree on {*u, v*} contains exactly one edge and displays {*u*} ∣ {*v*}. The conditional value 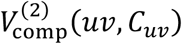 is defined only when that unique empirical edge carries a valid length. These maps are the direct one-edge specialization of the same reference/recovery logic, kept outside the frozen global domains so that neither *G* nor *B*^∗^ is changed.

*Proof: Appendix F*.*1*.

Figure 1D summarizes the layer boundary: the reference state is fixed by (*S, T*_*g*_); empirical topology refines only eligible cells; numerical values remain a separate conditional decoration.

## 5. A finite composite-coordinate universe

A fused path should not disappear into generic missingness. To compare such paths across loci, the present ledger records each by its exact set of primitive reference edges.

### 5.1 Member-set-keyed coordinates

Define

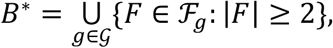

with the member set itself serving as the coordinate identifier.

#### Proposition 4 (finite composite-coordinate universe)

*B*^∗^ is finite and uniquely determined by (*S*, {*T*_*g*_}). Every *C* ∈ *B*^∗^ is a connected reference-tree path generated as a non-singleton fiber at least one locus.

*Proof: Appendix D*.*1*.

Within one locus, fibers are disjoint. Across loci, the same member set may be generated repeatedly and is deduplicated, while distinct elements of *B*^∗^ may overlap or nest. Thus *B*^∗^ is a ledger of possible composite coordinates, not an additive or orthogonal basis.

### 5.2 Fiber saturation

For *C* ∈ *B*^∗^, let

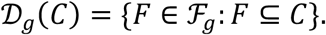

A coordinate is fiber-saturated when it contains no deleted edge and every current fiber intersecting *C* lies wholly inside *C*. Equivalently,

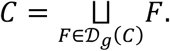

The equivalence follows because *F*_*g*_ partitions the nondeleted primitive edges. Fiber-saturated decomposability is the more general exact criterion. Requiring every primitive member to be eligible is more restrictive and defines only the subclass `primitive_decomposable`.

*Proofs of fiber saturation and unique whole-fiber decomposition are given in Appendices D*.*2–D*.*3*.

### 5.3 Composite reference states

Define

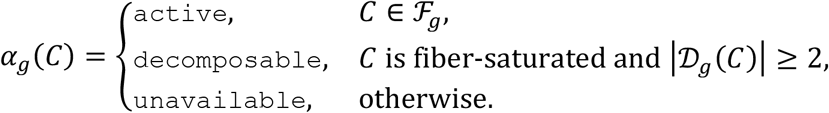

An active coordinate is one current fiber. A decomposable coordinate is the unique disjoint union of at least two whole current fibers. An unavailable coordinate has one or both of two defects: it contains a deleted primitive edge, or a current fiber intersects *C* but extends beyond *C*.

For example, if

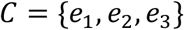

and the current fibers are {*e*_1_, *e*_2_} and {*e*_C_}, then *C* is decomposable into two whole fibers but is not `primitive_decomposable`.

*Proof: Appendix D*.*4*.

### 5.4 Component split families

For *F* ∈ *F*_*g*_, define

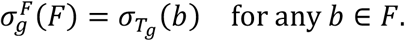

The direction same fiber ⇒ same projected split makes this definition well defined. The reverse direction same projected split ⇒ same fiber shows that distinct fibers have distinct projected splits, so 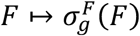 is injective.

For decomposable *C*, let

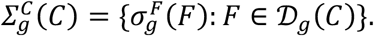

This is the split image of the unique member-set decomposition. It suffices for Boolean recovery but does not replace the member-set coordinate or imply empirical order or adjacency.

*Proof: Appendix D*.*5*.

### 5.5 Composite empirical recovery

Let 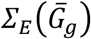 be the full edge-split set of the normalized empirical tree. Define

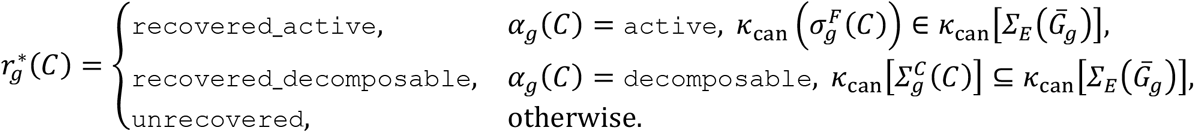

The state is total and single-valued. `recovered_decomposable` certifies only that the uniquely selected component splits are displayed; it does not imply adjacency, a continuous empirical path, or absence of intervening edges.

*Proof: Appendix D*.*6*.

### 5.6 Scope

The composite ledger is static and locuswise. It does not describe transformations across nested restrictions or representations; those questions lie outside the present study.

## 6. The finite categorical truth table and conditional numerical decoration

The primitive and composite constructions together define one finite categorical object. Here, categorical means taking values in finite state alphabets and invokes no category-theoretic structure. The term “truth table” denotes the complete finite assignment of those states under fixed labelled topological inputs and conventions; it is not a Boolean truth table, and the states are not Boolean-valued. Numerical estimates are attached only after the categorical states and their value-construction modes have been fixed.

### 6.1 Value-construction mode

For *C* ∈ *B*^∗^, define

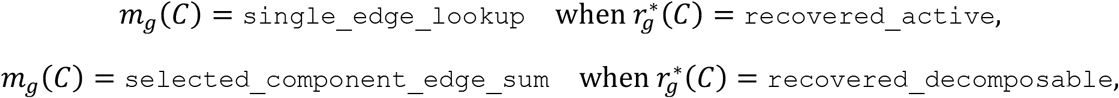

and

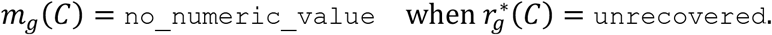

The mode *m*_*g*_ is categorical and topology-determined. Topology alone decides whether a later value, if lengths are supplied, is obtained by one edge lookup, by summing selected component edges, or not at all. The mode is a mathematical field of the ledger rather than a description of any particular software output, and it does not assert that lengths are present.

*Proof: Appendix E*.*1*.

### 6.2 The categorical truth table

Let

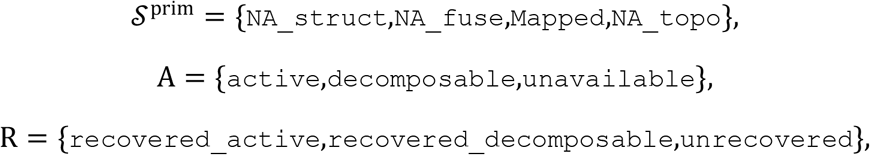

and

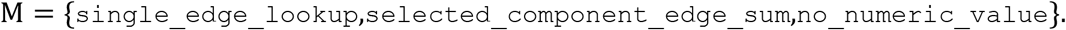

Define

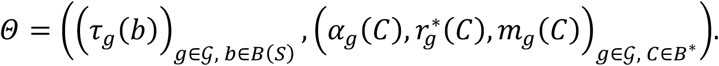

#### Theorem 5 (unique categorical truth table)

For fixed labelled topological inputs and fixed conventions, *Θ* is one uniquely determined element of

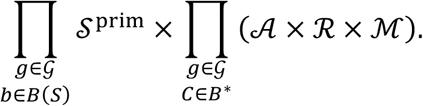

The index set and every alphabet are finite. “Unique” therefore means that the fixed inputs select one state for every cell. It does not mean that the chosen species tree is historically unique or correct, and it does not make an empirical branch-length estimate a truth state.

*Proof: Appendix E*.*2*.

### 6.3 Conditional numerical decoration

Assume that every normalized empirical tree carries a total nonnegative length function 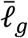. For a `Mapped` primitive coordinate *b*, let *e*_*b*_ be the unique empirical edge displaying 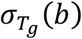. For recovered composites, let *e*_*c*_ be the unique edge displaying the active split and *e*_*F*_ the unique edge displaying the split of component fiber *F*.

Define

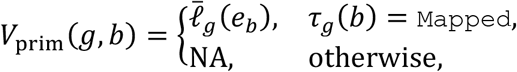

and

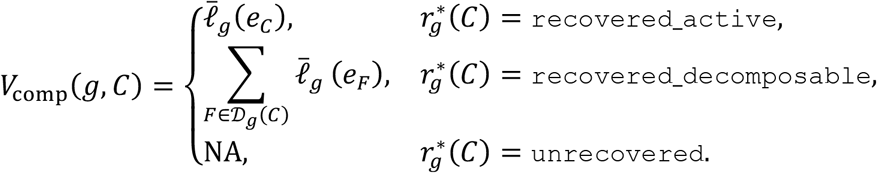

Collect these values as

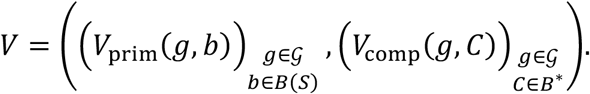

#### Corollary 6 (conditional numerical determinacy)

Conditional on the supplied normalized length functions, *V* is one single-valued element of a finite-index product with factors ℝ_≥0_ ⊔ {NA}.

The categorical theorem is the primary result; *V* is an infinite-valued decoration. A recovered-decomposable value is the sum of the uniquely selected component edges only. The selected edges need not be adjacent, need not form a continuous empirical path, and their sum is not asserted to be a full empirical path length, a decomposition of a recovered-active fused edge, or a sum of latent primitive member lengths. A recovered-active edge length is likewise never identified with latent primitive-member sums. Neither *Θ* nor *V* establishes historical evolutionary ground truth.

*Proof: Appendix E*.*3*.

## 7. Computational validation

The categorical results are mathematical statements proved in Appendices A-F. The computation reported here checks whether those definitions have been transcribed faithfully into executable state assignments on a finite domain; it is not a premise or proof of the theorem.

### 7.1 Validation design and methods

I performed a deterministic exhaustive validation on the six-taxon reference tree (A, (B,C), (D, (E, F))) ; The domain contained every retained subset of size three to six and every normalized labelled unrooted empirical topology on that subset, including multifurcations. A graph-free labelled set-partition recurrence supplied independent expected topology counts for each retained-set size, which were then compared with the topology generator.

For each retained set, the graph evaluator constructed the union of reference-tree paths, identified deleted original edges, and derived original-edge fibers by degree-two suppression. Every retained-taxon construction order was checked by passing its own ordered-anchor path-union edge set to an explicit-input fiber constructor and comparing both that edge set and the resulting fiber partition with the all-pairs construction.

The split evaluator independently projected each original-edge split to the retained taxa, treated empty projections as structural, and grouped equal nonempty projections. Primitive graph and split states were compared cellwise. For composite coordinates, the graph evaluator classified deletion and fiber saturation and queried graph-derived component splits. A separately implemented split evaluator classified structural contamination, partial or whole split-group coverage, and active, decomposable, or unavailable state; it then queried the empirical split index within its own recovery and numerical branches. The split evaluator did not call the graph composite state or recovery functions or consume their returned objects. The two evaluators shared the fixed coordinate and split representations and the common empirical topology and length input so that they answered the same cases.

For the certified two-taxon boundary, I explicitly constructed the normalized empirical tree consisting of two labelled leaves joined by one edge, built its empirical split index, and assigned that edge the declared valid length 1. Separate graph- and split-side boundary functions derived and queried the corresponding split. The one-taxon boundary was evaluated separately by confirming an empty retained-edge ledger and structural state for every frozen primitive coordinate. Main-domain empirical edge lengths were deterministic nonnegative decimal values assigned by canonical split rank; active coordinates used a single-edge lookup and recovered decomposable coordinates used the sum of selected component-edge lengths.

Complete case, primitive, composite, order, boundary, recurrence, coverage, invariant, and discrepancy tables were persisted. Two clean runs were compared bytewise. A separate output-only QA script did not import the validation implementation, and an abstract-syntax-tree audit checked prohibited call boundaries between the compared evaluators.

### 7.2 Validation results

The exhaustive domain contained 42 retained-taxon sets and 472 normalized empirical-tree cases: 260 binary and 212 multifurcating. These cases produced 4,248 primitive cells and 9,912 composite cells on the frozen 21-coordinate composite axis, for 14,160 main categorical cells in total (Table 1).

**Table 1.** Summary of the exhaustive computational validation.

| Validation component | Enumerated scope | Result |
| --- | --- | --- |
| Normalized empirical-tree domain | 472 cases: 260 binary; 212 multifurcating | Complete |
| Primitive graph/split comparison | 4,248 cells | 4,248 agreements |
| Separately implemented composite comparison | 9,912 cells | 9,912 agreements |
| Construction and suppression order | 1,920 and 84 trials | All passed |
| Explicit two-leaf empirical lookup | 15 cases | All passed |
| One-taxon boundary | 6 cases | All passed |
| Independent topology-count recurrence | Retained-set sizes 3-6 | 1, 4, 26, 236; generator matched |
| Reproducibility and QA | 13 files; 40 output checks; 6 static checks | All identical or passed |
| Machine-readable discrepancies | All comparisons | 0 data rows |

The graph and split constructions agreed in all 4,248 primitive cells. The separately implemented graph- and split-side composite evaluators agreed in all 9,912 composite cells on reference state, saturation, component member sets, component splits, recovery state, construction mode, and conditional numerical value. The machine-readable discrepancy table contained its required header and zero data rows.

All 1,920 construction-order trials agreed on both the ordered-anchor path-union edge set and the fibers derived from that observed edge set; all 84 suppression-order trials also passed. All 15 explicit normalized two-leaf empirical lookups agreed between the graph and split boundary functions and returned `recovered_active`, `single_edge_lookup`, and the declared value 1. All six one-taxon boundary cases assigned structural state to every primitive coordinate and invoked no empirical recovery step. The independent set-partition recurrence and topology generator both produced 1, 4, 26, and 236 topologies for three through six retained taxa.

Primitive output totals were 2,406 `Mapped`, 890 `NA_topo`, 598 `NA_fuse`, and 354 `NA_struct`. Composite reference totals were 280 active, 6,912 decomposable, and 2,720 unavailable; compositerecovery totals were 221 `recovered_active`, 459 `recovered_decomposable`, and 9,232 unrecovered.

Two clean executions produced byte-identical hashes for all 13 canonical files. Output-only QA passed 40 of 40 checks, and the static call-boundary audit passed 6 of 6 checks. These results support implementation fidelity on the enumerated finite domain; they do not constitute a computational proof of the general theorem or certification of production software.

## 8. Discussion

This study addresses a representational premise that precedes branch-level evolutionary inference: whether a branch coordinate is actually the same object across loci. The answer cannot be obtained from a branch label alone when taxa differ among loci. The underlying restriction and split-display operations are standard. The contribution is to retain the original-edge provenance normally discarded by reduction and assemble it into a cross-locus ledger. For each locus, the pointed map records every frozen reference edge as deleted or as a member of the complete contraction fiber of one reduced edge, making the structural, fused, and eligible states explicit before empirical recovery.

Projected splits faithfully encode these graph fibers. This fiber–split interface follows from standard restriction and edge–split uniqueness; its role here is to connect the provenance ledger to a practical split representation, not to introduce an alternative definition of branch identity. Two surviving reference edges have the same projected split exactly when they belong to the same fiber. The equivalence also clarifies why empty-side cases must be separated as structural deletion rather than pooled with ordinary split matching.

Earlier split-based concordance methods had already encountered the ambiguity produced by incomplete taxon sampling. PhyParts compares rooted gene-tree clades with a reference species tree and permits one input clade to be consistent with several reference clades; its implementation assigns the concordant signal to the most nested (shallowest) consistent species-tree node, while conflicting signal is grouped and summarized through node-level concordance, conflict, and internode-certainty (ICA) statistics (Smith et al. 2015). This treatment is appropriate for its objective of summarizing phylogenetic signal, but it does not retain the multiple consistent reference edges as the provenance of a new coordinate. From the coordinate-ledger perspective developed here, the multi-match is not merely uncertainty about which primitive branch should receive the observation. It is the split signature of a restriction-induced fiber in which those primitive edges have ceased to be separately identifiable and survive instead as one composite coordinate.

SplitAligner made this representational step operational by retaining the complete original-edge member set of each multi-match, assigning fusion states to its primitive members, and carrying the resulting composite coordinate into a cross-locus matrix. The present theorem supplies the graph-theoretic explanation: degree-two suppression uniquely determines the fiber, and fiber–split equivalence proves that the shared projected split is a faithful encoding of that fused coordinate rather than an arbitrary ambiguity-resolution rule.

Related combinatorial traditions reconstruct trees from compatible full splits and represent collections of partial taxon splits with Buneman graphs, weak X-trees, and subtree distances. These are adjacent but not equivalent objects. Their coordinates are taxon splits, whereas the composite coordinates here are sets of frozen reference edges retained as restriction provenance; they do not supply the locus-specific active, decomposable, unavailable, and empirical-recovery states assembled in this ledger (Buneman 1971; Semple and Steel 2003; Bryant et al. 2025).

The distinction between reference truth and empirical recovery is equally important. Structural deletion and fusion are determined before an empirical gene tree is examined. A discordant empirical topology cannot convert a structural coordinate into a fused one, split a fused fiber back into primitive members, or make a deleted reference edge eligible. It can only determine whether an eligible split is recovered. The four-state primitive output is therefore a composition of a three-state graph partition with a two-state refinement of one partition block.

This separation limits the meaning of `Mapped` and `NA_topo`. `Mapped` means that the normalized empirical tree displays the reference coordinate’s restricted split. It does not mean that the reference branch is historically correct, that the empirical gene tree is error free, or that the branch length is an unbiased estimate. Conversely, `NA_topo` records absence of the eligible reference split from the supplied empirical topology. It does not identify whether the discordance arose from incomplete lineage sorting, introgression, gene-tree estimation error, or another process.

An unresolved reference tree and an unresolved empirical tree therefore have different meanings in this framework. At a reference polytomy, an unchosen refinement has no primitive coordinate and lies outside the ledger. At an empirical polytomy, the eligible reference coordinate still exists, but its split may be absent from the normalized empirical tree and is recorded coarsely as `NA_topo`. The present theorem does not subdivide `NA_topo` according to why the split is absent, and it does not prescribe arbitrary resolution of a biological polytomy.

Terminal branches provide a useful internal check. A retained taxon is represented by a terminal edge in every empirical tree on that taxon set. Hence a genuine retained terminal coordinate can be `Mapped` or fused, but not structural or `NA_topo`. An internal reference edge that projects to the same singleton split is not confused with that terminal coordinate, because endpoint collapse places it in the terminal edge’s non-singleton fiber. The full edge-split set, including terminal splits, is therefore required for a correct recovery theorem.

Composite coordinates make explicit what is otherwise hidden by a primitive-only matrix. When several primitive edges fuse, their member rows are correctly marked `NA_fuse` because their individual identities are unavailable. At the same time, the non-singleton fiber survives as an active composite coordinate. Across loci, the resulting composite ledger may contain overlapping or nested member sets. This does not make the ledger ambiguous, but it does mean that the composite universe is not a basis whose columns can be added or compared without regard to locus-specific representation. The ledger does not prescribe simultaneous use of overlapping composites as independent predictors or matrix columns; any downstream analysis must declare a representation rule and must not treat the composite universe as an additive or orthogonal basis.

Fiber saturation governs when a previously generated composite can be represented at another locus. A composite is active if it is one current fiber. It is decomposable if it is exactly the union of a uniquely selected set of current fibers. A candidate that cuts through a current fiber is unavailable. This rule prevents an analyst from selecting only part of an indivisible current coordinate.

Recovered decomposable coordinates require especially careful interpretation. The selected empirical edges are identified by the splits of the current fiber components. They need not form an adjacent chain in the empirical tree. Their sum is a declared construction over selected components, not a newly inferred empirical path. It is also not a claim that a fused empirical edge can be decomposed into latent primitive contributions. The categorical construction mode is therefore retained as an explicit field rather than hidden inside a number.

The separation of *Θ* and *V* expresses the main conceptual boundary of the paper. The categorical object is length free. It states which coordinates are structural, fused, eligible, recovered, decomposable, or unavailable, and how a recovered numerical value would be constructed. Numerical values enter only after valid empirical lengths are supplied. Their single-valuedness is conditional on those inputs and on edge–split injectivity in the normalized tree.

The two-taxon boundary illustrates the same logic in its simplest form. Restriction to a taxon pair reduces the reference tree to the unique path connecting the pair. Every member of that path fuses into one composite coordinate, while all off-path edges are structural. The empirical two-leaf tree contains one edge, so the active composite is recovered by one edge lookup. The pairwise scalar may therefore be annotated with deterministic path membership. Path membership alone, however, does not assign numerical contributions to the path’s primitive edges.

The finite validation provides a computational check on the translation from the definitions to executable state assignments. Agreement between graph- and split-derived evaluators across all enumerated six-taxon cases supports implementation fidelity on that domain, including multifurcations and the two-taxon boundary. It does not extend the proof beyond its stated assumptions or certify production software.

The present theory intentionally stops before several broader questions. It does not characterize how fibers transform under nested taxon restrictions, prove a no-unfusion law across restrictions, or establish a general representation-factorization theorem. It also does not quantify downstream errors caused by assigning a valid composite value to an invalid primitive address. These are related but distinct problems.

The practical implication is that branch-resolved comparative analyses should expose their coordinate ledger before interpreting their numerical matrix. A missing primitive value may indicate structural deletion, fusion into a composite coordinate, or topological non-recovery. These mechanisms have different meanings and should not be collapsed merely because they share an NA representation in a table.

More broadly, the result identifies branch identity as a graph-theoretic object conditional on a reference tree and a taxon universe. Once those inputs are fixed, the categorical ledger is not a matter of software preference. It is uniquely determined.

## Supporting information

Supplementary Computational Evidence

## 9. Data and code availability

No new empirical biological data were generated. All definitions, theorem statements, and proofs supporting the mathematical results are contained in this manuscript. The computational validation scripts, frozen protocol addendum, complete machine-readable result tables, discrepancy ledger, execution logs, clean-rerun hash comparisons, output-only QA, and static call-boundary audit are supplied with the revised preprint in the versioned Supplementary Computational Evidence Package, `BranchIdentity_Computational_Evidence_v0.2_20260723_JST.zip`. The computation evaluates implementation fidelity on the enumerated six-taxon domain and is not a premise of any proof. Production SplitAligner software and application-specific data remain associated with the companion study (Wu 2026).

## 10. Scope statement

This study establishes:

- the canonical pointed restriction ledger on the frozen reference edge axis;
- the structural, fused, and eligible primitive classification;
- the fiber–split interface and eligible-only empirical refinement;
- the member-set-keyed composite-coordinate universe and fiber-saturation rule;
- the total composite reference and empirical recovery states;
- the finite categorical truth table and its fixed-input cellwise determinacy;
- conditional numerical determinacy under supplied normalized lengths;
- the two-taxon and one-taxon boundaries.

This study does not establish:

- historical correctness of a species tree or gene tree;
- unbiasedness or biological truth of empirical branch lengths;
- latent primitive decomposition of an empirical fused edge;
- nested-restriction transformation laws;
- general representation invariance;
- downstream statistical consequences of coordinate misuse;
- empirical performance of production software.

These boundaries are part of the theorem statement, not merely limitations added after the result.

## APPENDIX A. Preliminary tree results and empirical normalization

All notation and standing assumptions are as in Section 2. The results collected here establish the elementary tree facts used in the later proofs and the uniqueness of the empirical-tree normalization.

### A.1 Basic tree facts

#### Lemma A1 (unique paths and edge chains)

In a finite tree, every two vertices are joined by exactly one simple path. Consequently, every two edges are joined by a unique minimal edge chain.

**Proof**. Connectedness guarantees the existence of a path between any two vertices. If two distinct simple paths joined the same pair of vertices, their union would contain a cycle, contradicting acyclicity.

For edges *e* and *f*, subdivide each tree edge once and let *m*_*e*_ and *m*_*f*_ denote the subdivision vertices corresponding to e and *f*. The subdivided graph is again a tree, so it contains a unique simple path from *m*_*e*_ to *m*_*f*_. Reading the original edges encountered along this path gives a unique consecutive edge sequence

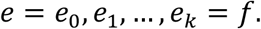

This sequence is minimal because removing either endpoint edge would disconnect the chain from *e* or *f*.

#### Lemma A2 (minimal connecting subtree and two-sidedness)

For every nonempty retained-taxon set *T*_*g*_ ⊆ *T*, there is a unique minimal subtree *K*_*g*_ ⊆ *S* connecting *T*_*g*_. It is the union of the unique paths in *S* between retained taxa. Every degree-one vertex of *K*_*g*_ belongs to *T*_*g*_, and for every edge *e* ∈ *E*=(*K*_*g*_ ), both components of *K*_*g*_ − *e* contain at least one retained taxon.

**Proof**. Fix *t*_0_ ∈ *T*_*g*_ and let *U*_*g*_ be the union of the unique paths in *S* from *t*_0_ to all other members of *T*_*g*_. The subgraph *U*_*g*_ is connected and contains *T*_*g*_. The unique path between any two retained taxa is contained in the union of their paths to *t*_0_, so *U*_*g*_ is also the union of all pairwise retained-taxon paths.

Every connected subgraph of *S* containing *T*_*g*_ must contain the unique path between each retained pair and must therefore contain *U*_*g*_. Hence *U*_*g*_ is the unique inclusion-minimal connected subgraph containing *T*_*g*_. Set *K*_*g*_ = *U*_*g*_

If a degree-one vertex of *K*_*g*_ were not retained, deleting that vertex and its incident edge would leave a smaller connected subgraph containing *T*_*g*_, contradicting minimality. Similarly, if one component of *K*_*g*_ − *e* contained no retained taxon, that component and the edge *e* could be removed without disconnecting the retained taxa. This would again contradict minimality.

#### Lemma A3 (finiteness)

The sets *B*(*S*), *E*(*K*_*g*_), *F*_*g*_, *B*^∗^, and all primitive and composite cell-index sets used in the manuscript are finite.

**Proof**. The reference tree *S* is finite and therefore has finitely many edges. Each *K*_*g*_ is a subgraph of *S*, and each *F*_*g*_ is a partition of the finite set *E* (*K*_*g*_). The locus set *G* is finite, so the union of all non-singleton fibers generated across loci is finite. The remaining cell-index sets are finite Cartesian products of these finite sets.

### A.2 Canonical normalization of empirical trees

#### Lemma A4 (canonical empirical-tree normalization)

Let *G* be an admissible raw empirical tree with labelled leaf set *T*_*g*_, after root orientation has been forgotten when applicable. Join two edges of *G* in an auxiliary graph *H*_2_(*G*) exactly when they meet at a degree-two vertex of *G*. Contract every connected component of *H*_2_ (*G*) to one edge. Then:

1. the components of *H*_2_(*G*) are the uniquely determined maximal degree-two edge chains of *G*;
2. the result of contraction is independent of suppression order;
3. the normalized tree 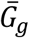 is unique up to leaf-label-preserving unrooted isomorphism;
4. 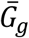has labelled leaf set *T*_*g*_ and no degree-two internal vertex; and
5. if *G* carries a total finite nonnegative edge-length function, assigning each quotient edge the sum of the lengths on its contracted chain defines a unique normalized length function 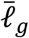.

**Proof**. The connected components of *H*_2_(*G*) uniquely partition *E*(*G*). At each endpoint of a tree edge, adjacency in *H*_2_(*G*) is possible only when that endpoint has degree two, in which case there is exactly one other incident tree edge. Every vertex of *H*_2_(*G*) therefore has degree at most two.

A cycle in *H*_2_(*G*) would correspond to a closed sequence of distinct consecutive edges in *G*, producing a cycle in *G*. Since *G* is acyclic, every component of *H*_2_(*G*) is a simple path or a singleton. Its corresponding tree edges form a consecutive chain whose internal junctions all have degree two. Component maximality shows that this chain is maximal.

The chain partition is fixed before any contraction is performed. Sequential suppression in any order merely contracts subsets of the same components and therefore yields the same quotient incidence relation. Contracting paths in a tree cannot create a loop or parallel edges, because either would lift to a cycle in the original graph.

The endpoints of a maximal chain have degree one or degree different from two. Consequently, the quotient has no degree-two internal vertex. Labelled leaves are never internal junctions of the contracted chains, so their labels and degree-one status are retained. The admissibility condition on rooted inputs excludes the creation of an unlabelled leaf when root orientation is forgotten. The quotient is therefore uniquely determined up to leaf-label-preserving unrooted isomorphism.

If lengths are supplied, every quotient edge corresponds to one unique finite chain *Q* ⊆ *E*(*G*). Define

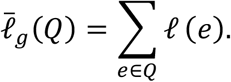

Because *Q* is uniquely determined and finite addition is associative and commutative, the value does not depend on the order of suppression.

The categorical conclusions of Lemma A4 require no branch lengths. Its final conclusion is conditional on a total valid length function on the raw empirical tree.

## APPENDIX B. Reference restriction and primitive-coordinate states

### B.1 Proof of Proposition 1

#### Proof of Proposition 1 (canonical pointed restriction ledger)

By Lemma A2, the minimal subtree *K*_*g*_ connecting *T*_*g*_ exists and is unique. It follows immediately that

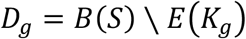

is uniquely determined.

The auxiliary graph *H*_*g*_ is determined entirely by the incidences and vertex degrees in *K*_*g*_: its vertices are the edges of *K*_*g*_, and two are adjacent exactly when they meet at a degree-two vertex of *K*_*g*_. Its connected components therefore form a unique partition *F*_*g*_ of *E* (*K*_*g*_).

As in the proof of Lemma A4, every vertex of *H*_*g*_ has degree at most two, and a cycle in *H*_*g*_ would induce a cycle in *K*_*g*_. Each component is consequently a simple path or a singleton. The corresponding reference edges form a maximal consecutive chain whose internal junctions have degree two in *K*_*g*_. Every fiber is thus a connected path segment of *K*_*g*_, and hence of *S*.

Finally, every primitive edge *b* ∈ *B*(*S*) satisfies exactly one of two alternatives. If *b* ∈ *D*_*g*_, then *π*_*g*_(*b*) =⊥. Otherwise *b* ∈ *E* (*K*_*g*_) and belongs to exactly one component of *H*_*g*_, to which *π*_*g*_ assigns it. Thus

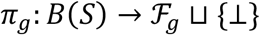

is total and single-valued. All objects in the pointed restriction ledger are therefore uniquely determined by (*S, T*_*g*_).

The proof uses no empirical topology, split canonicalization, or branch length. The binary assumption on *S* is not required for the restriction construction itself.

### B.2 Proof of Lemma 1

#### Proof of Lemma 1 (fiber–split interface)

Let *b, b*′ ∈ *E* (*K*_*g*_). By Lemma A2, both components created by deleting either edge contain retained taxa. Hence 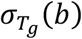 and 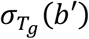 are both nondegenerate unordered splits of *T*_*g*_.

First suppose that

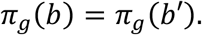

Then *b* and *b*′ belong to the same component of *H*_*g*_. There is a sequence

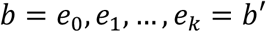

in which each consecutive pair meets at a degree-two vertex of *K*_*g*_. At such a junction there is no third incident edge and therefore no off-chain component containing a retained taxon. Moving a cut from *e*_*i* − 1_ to *e*_*i*_ crosses no retained branch and leaves the induced unordered partition of *T*_*g*_ unchanged. Repeating this argument along the chain gives

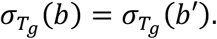

Conversely, suppose that

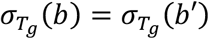

but *b* and *b*′ belong to different fibers. Let

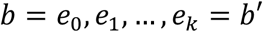

be their unique minimal edge chain in *K*_*g*_, given by Lemma A1. Because the two edges are not in the same component of *H*_*g*_, at least one junction *v* = e_*i* − 1_ ∩ e_*i*_ on this chain has degree at least three in *K*_*g*_. Choose an edge incident to *v* that does not belong to the joining chain. Following this edge away from *v* eventually reaches a degree-one vertex of *K*_*g*_, which is a retained taxon by Lemma A2; call it *x*.

Again by Lemma A2, choose a retained taxon *p* in the component of *K*_*g*_ − *b* not containing *b*′, and a retained taxon *q* in the component of *K*_*g*_ − *b*′ not containing *b*. Under the cut at *b*, the off-chain taxon *x* lies on the same side as *q* and opposite *p*. Under the cut at *b*′, it lies on the same side as *p* and opposite *q*. The two unordered splits are therefore different, contradicting the assumption.

Hence

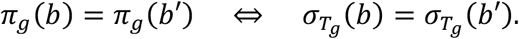

The fibers are defined by graph restriction; the split equality is a faithful encoding of those already determined fibers.

### B.3 Primitive reference trichotomy

#### Proposition 2 (primitive reference trichotomy)

For every *g* ∈ *G*and *b* ∈ *B*(*S*), exactly one of the following applies:

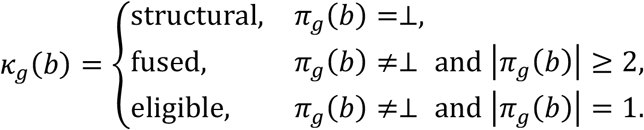

At a fixed locus, every fused primitive edge belongs to exactly one non-singleton active fiber.

**Proof**. Proposition 1 establishes that *π*_*g*_ is total and single-valued. For any *b* ∈ *B*(*S*), either *π*_*g*_(*b*) =⊥, or *π*_*g*_(*b*) is a member of the finite partition *F*_*g*_. In the second case, its cardinality is either one or at least two. These alternatives are mutually exclusive and exhaustive, so *K*_*g*_ is total and single-valued.

If *b* is fused, the unique fiber *π*_*g*_(*b*) has at least two members. Because *F*_*g*_ is a partition, no second fiber at the same locus can contain *b*. Thus every fused primitive edge belongs to exactly one active non-singleton fiber at that locus. No uniqueness across different loci is implied.

### B.4 Endpoint collapse

#### Corollary B1 (endpoint collapse)

Let *x* ∈ *T*_*g*_, let *b*_*x*_ be the terminal reference edge incident to *x*, and let *b* ≠ *b*_*x*_ be an internal reference edge satisfying

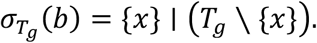

Then

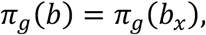

their common fiber contains at least two edges, and both *b* and *b*_*x*_ are fused.

**Proof**. The terminal edge *b*_*x*_ induces the same singleton split

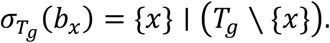

Both edges are nondeleted because both sides of this split are nonempty. Lemma 1 therefore gives

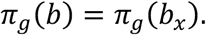

Since *b* ≠ *b*_*x*_, their common fiber has at least two members. Proposition 2 classifies both edges as fused.

## APPENDIX C. Empirical recovery and unique edge addresses

### C.1 Proof of Lemma 2

#### Proof of Lemma 2 (edge–split injectivity)

Let *G* be a finite, simple, connected, acyclic, unrooted tree whose degree-one vertices are bijectively labelled by a taxon set *X* and whose internal vertices do not have degree two. Take distinct edges *e* ≠ *f* and let

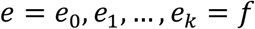

be their unique minimal edge chain, as supplied by Lemma A1. Since *e* ≠ *f, k* ≥ 1, and each consecutive pair meets at a junction vertex *v*_*i*_ = *e*_*i* − 1_ ∩ *e*_*i*_.

Assume for contradiction that

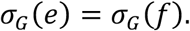

If some junction *v*_*i*_ had degree at least three, choose an incident edge that is not on the joining chain. Following a nonbacktracking path away from *v*_*i*_ through that edge must terminate, by finiteness and acyclicity, at a degree-one vertex. That vertex has a taxon label by hypothesis; call it *x*. Likewise, choose a labelled leaf *p* in the component of *G* − *e* away from *f*, and a labelled leaf *q* in the component of *G* − *f* away from e. Such leaves exist because each finite component obtained by cutting a tree contains a degree-one vertex of the original tree.

At the cut e, the leaves *x* and *q* lie on the same side and opposite *p*. At the cut *f*, the leaves *x* and *p* lie on the same side and opposite *q*. The two labelled splits are therefore different, contradicting the assumption. Hence every junction *v*_*i*_ on the joining chain must have degree two. Each such junction is internal, contrary to the normalization assumption. Thus distinct edges display distinct unordered labelled splits.

The no-degree-two condition is necessary for this theorem class: the two edges incident to an internal degree-two vertex display the same labelled split. Full leaf labelling is also needed for the general statement, because an off-chain component ending only in an unlabelled leaf need not distinguish the induced splits of the labelled taxa. Binary resolution is not required; an internal degree of at least three supplies the off-chain labelled component used in the proof.

### C.2 Proof of Proposition 3

#### Proof of Proposition 3 (eligible-only empirical refinement)

Fix *g* ∈ *G*. For every eligible primitive coordinate *b*, the projected split 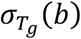 is fixed by (*S, T*_*g*_). Lemma A4 fixes the normalized empirical tree 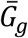 up to leaf-label-preserving unrooted isomorphism and therefore fixes its finite edge-split set 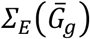.

The faithful canonicalization *K*_*can*_ satisfies

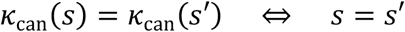

for unordered labelled splits. Consequently,

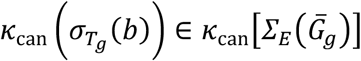

has exactly one Boolean truth value. The eligible-axis recovery map *r*_*g*_ is therefore total and single-valued.

Proposition 2 partitions the primitive reference axis into the disjoint structural, fused, and eligible blocks. Assigning `NA_struct` to the structural block, `NA_fuse` to the fused block, and refining only the eligible block by *r*_*g*_ ∈ {`Mapped`,`NA_topo`} produces a total and single-valued map

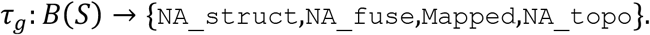

A leaf-label-preserving unrooted isomorphism preserves every displayed labelled split and hence preserves *r*_*g*_ and τ_*g*_. No invariance under arbitrary leaf relabelling is claimed. Finally, *K*_*g*_, *D*_*g*_, *H*_*g*_, *F*_*g*_, *π*_*g*_, and *K*_*g*_ were defined before and independently of 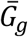. Empirical recovery therefore cannot alter the reference restriction ledger.

The categorical membership test itself does not require edge–split injectivity. Injectivity becomes necessary only when a displayed split must be assigned a unique empirical edge address for numerical lookup.

### C.3 Retained terminal coordinates

#### Corollary C1 (retained-terminal dichotomy)

Let *x* ∈ *T*_*g*_ and let *b*_*x*_ be the terminal reference edge incident to *x*. Then

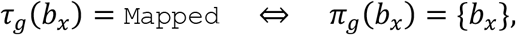

and

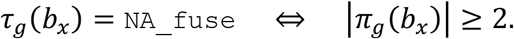

Consequently, a retained terminal coordinate is never `NA_struct` or `NA_topo`.

**Proof**. Because *x* is retained, the terminal edge *b*_*x*_ belongs to the minimal connecting subtree *K*_*g*_ and is not structural. If its reference fiber is non-singleton, Proposition 2 classifies it as fused, so the combined primitive state is `NA_fuse`.

If *π*_*g*_(*b*_*x*_) = {*b*_*x*_}, then *b*_*x*_ is eligible. Every empirical tree with leaf set *T*_*g*_, including 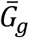, contains a terminal edge incident to *x* and displays the singleton split

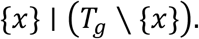

Because empirical recovery uses the full edge-split set, including terminal splits, the eligible terminal coordinate is `Mapped`. The singleton and non-singleton fiber cases are exhaustive, proving both biconditionals and the exclusion of `NA_struct` and `NA_topo`.

Corollary B1 explains the non-singleton case: when an internal reference edge projects to the same singleton split, it and *b*_*x*_ belong to one endpoint-collapse fiber and are jointly fused.

### C.4 Conditional paired-design bookkeeping

Let ℰ_*g*_ be an explicitly stated paired-evidence domain and let

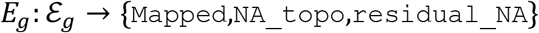

be a fixed total deterministic map. The arguments supplied to *E*_*g*_ are already single-valued by Propositions 1–3 and the associated recovery definitions. Evaluating a total single-valued function at fixed arguments gives exactly one output. The paired assignment is therefore single-valued.

This conclusion is conditional on the declared map *E*_*g*_. A different total deterministic evidence rule may produce a different, but internally deterministic, bookkeeping assignment. Because *E*_*g*_ is external paired-design bookkeeping, `residual_NA` is not a fifth graph-theoretic mechanism and is not an additional state in the categorical truth table *Θ*.

## APPENDIX D. Composite-coordinate existence, representation, and recovery

### D.1 Proof of Proposition 4

#### Proof of Proposition 4 (finite composite-coordinate universe)

For each *g* ∈ *G*, Proposition 1 uniquely determines the finite partition *F*_*g*_ and hence its collection of non-singleton fibers. Define

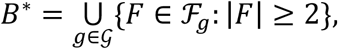

with equality keyed by the exact member subset of the fixed primitive edge set *B*(*S*). Because *G*is finite and each *F*_*g*_ is finite, *B*^∗^ is finite. Every member of *B*^∗^ is a connected reference-tree path generated as a non-singleton restriction fiber at least one locus.

Within one locus, distinct fibers are disjoint because they are blocks of one partition. Across loci, no common partition is asserted, so two elements of *B*^∗^ may overlap or be nested. These set-theoretic facts provide no addition, linear independence, or orthogonality. Thus *B*^∗^ is a member-set-keyed coordinate ledger, not an additive or orthogonal basis.

#### D.2 Fiber saturation

For *C* ∈ *B*^∗^, define

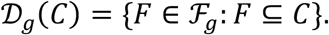

#### Lemma D1 (fiber-saturation equivalence)

For every nonempty *C* ⊆ *B*(*S*), the following are equivalent:

1. *C* ∩ *D*_*g*_ = Ø and every current fiber intersecting *C* is wholly contained in *C*;
2. *C* is the disjoint union of the whole current fibers contained in it:

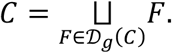

**Proof**. Suppose the first condition holds. Every *b* ∈ *C* is nondeleted and therefore lies in one unique partition block *F*_I_ ∈ *F*_*g*_. Since *F*_I_ ∩ *C* ≠ Ø, fiber saturation gives *F*_I_ ⊆ *C*, so *F*_I_ ∈ *D*_*g*_(*C*). The union of *D*_*g*_(*C*) therefore covers *C*, while every included fiber is by definition contained in *C*. The union is disjoint because *F*_*g*_ is a partition.

Conversely, suppose the displayed disjoint-union identity holds. Then *C* consists entirely of ordinary fiber members and contains no deleted edge. If a current fiber *F* intersects *C*, it intersects one block in the displayed union. Distinct blocks of the partition are disjoint, so *F* equals that included block and is wholly contained in *C*. Thus *C* is fiber-saturated.

### D.3 Unique whole-fiber decomposition

#### Lemma D2 (unique whole-fiber decomposition)

Whenever *C* is fiber-saturated, *D*_*g*_(*C*) is its unique decomposition into whole current fibers.

**Proof**. The family *D*_*g*_(*C*) is fixed by set comprehension from the fixed partition *F*_*g*_. Every *b* ∈ *C* lies in one unique block of that partition. Any union of whole partition blocks equal to *C* must therefore contain exactly the blocks containing the members of *C*, namely the elements of *D*_*g*_(*C*). No alternative whole-fiber decomposition exists.

### D.4 Composite reference states

#### Proposition D1 (composite reference trichotomy)

For *g* ∈ *G*and *C* ∈ *B*^∗^, define the composite reference state by the following mutually exclusive clauses:

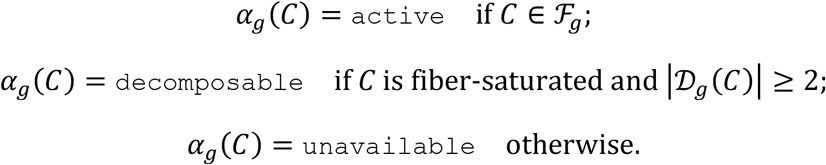

Then active, decomposable, and unavailable are mutually exclusive and exhaustive. More precisely:

1. *C* is active exactly when it is fiber-saturated with one component fiber;
2. *C* is decomposable exactly when it is the unique disjoint union of at least two whole current fibers; and
3. *C* is unavailable exactly when it contains a deleted primitive edge or cuts through a current fiber, that is, when

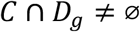

or

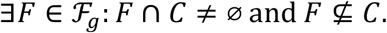

**Proof**. By Lemma D1, every nonempty fiber-saturated *C* has at least one component fiber. If it has one component, its disjoint union equals that fiber, so *C* ∈ *F*_*g*_ and is active. Conversely, an ordinary fiber is disjoint from *D*_*g*_, intersects no other fiber, and is saturated with itself as its unique component.

If a saturated *C* has at least two components, Lemma D2 gives its unique decomposition into at least two whole fibers, so it is decomposable. Conversely, such a whole-fiber union is saturated by Lemma D1. Active and decomposable are disjoint because their component counts are one and at least two. Every saturated coordinate falls into exactly one of these two cases.

The remaining coordinates are precisely the nonsaturated ones. Negating the saturation condition yields either a deleted member or a current fiber that intersects but is not contained in *C*, which is exactly the displayed unavailable criterion. Thus the three states are mutually exclusive and exhaustive.

The optional label `primitive_decomposable` is a subclass of decomposable obtained by additionally requiring every component fiber in *D*_*g*_(*C*) to be a singleton. It is not the general exact-representability criterion.

### D.5 Component split families

For *F* ∈ *F*_*g*_, define

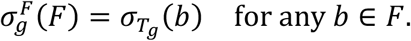

#### Lemma D3 (well-defined and injective fiber-split map)

The map 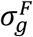 is well-defined and injective on *F*_*g*_.

**Proof**. All primitive edges in one fiber have the same projected split by Lemma 1, so the definition does not depend on the selected member *b* ∈ *F*. If 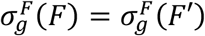, choose *b* ∈ *F* and *b*′ ∈ *F*′. Lemma 1 gives π_*g*_(*b*) = π_*g*_(*b*′), hence *F* = *F*′.

For decomposable *C*, define its component split family by

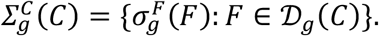

Because *D*_*g*_(*C*) is unique and 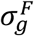 is injective, this set is the injective split image of the unique member-set decomposition, with no multiplicity loss. It is sufficient for the Boolean recovery question of whether all component splits are displayed. It does not replace the member-set coordinate, and it asserts no order, adjacency, or continuous empirical path among the realizing edges.

### D.6 Composite empirical recovery

#### Proposition D2 (composite empirical recovery)

Let 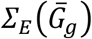 be the full edge-split set of the normalized empirical tree and define the recovery state by the following clauses:

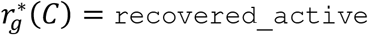

when α_*g*_(*C*) = `active` and

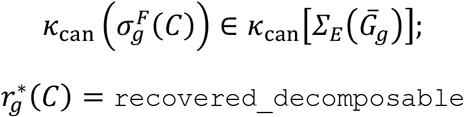

when α_*g*_(*C*) = `decomposable` and_*g*_

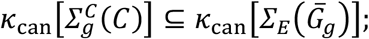

and set

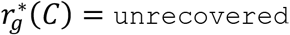

otherwise.

Then 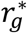 is total and single-valued.

**Proof**. Proposition D1 assigns exactly one composite reference state, so the active and decomposable recovery branches are mutually exclusive. In the active branch, membership of one fixed split in the fixed empirical split set has one Boolean truth value. In the decomposable branch, inclusion of one fixed finite split family in the fixed empirical split set likewise has one Boolean truth value. If the relevant condition fails, or if *C* is unavailable, the exhaustive otherwise branch assigns unrecovered. Exactly one recovery state is therefore assigned to every pair (*g, C*).

For a decomposable coordinate, `recovered_decomposable` certifies only that all uniquely selected component splits are displayed. It does not imply that their empirical edges are adjacent, form a path, or reproduce one fused empirical edge.

## APPENDIX E. Assembly of the categorical matrix and conditional numerical decoration

### E.1 Value-construction mode

For *C* ∈ *B*^∗^, define

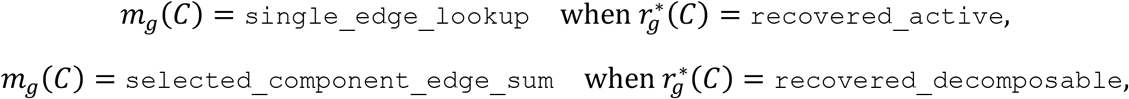

and

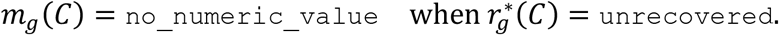

#### Lemma E1 (determinacy of construction mode)

The map *m*_*g*_ is total and single-valued and is defined without branch lengths.

**Proof**. Proposition D2 gives a total three-state partition for 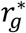. The displayed case map assigns one and only one construction mode to each recovery state. Its definition reads only the categorical recovery state, so it exists even when no normalized length function is supplied.

### E.2 Proof of Theorem 5

#### Proof of Theorem 5 (unique categorical truth table)

Let

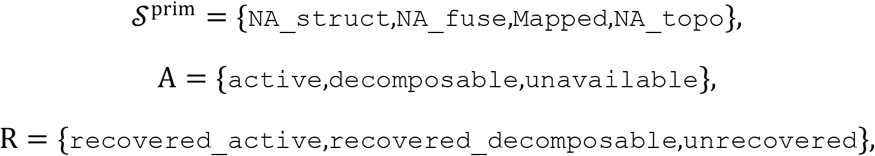

and

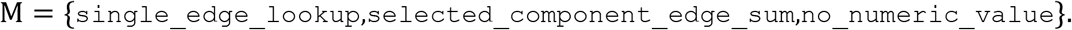

These four alphabets are finite. Lemma A3 establishes that *G, B*(*S*), and *B*^∗^ are finite.

For each primitive cell (*g, b*), Proposition 1 and Proposition 2 determine exactly one reference state, and Proposition 3 refines only the eligible case to determine exactly one value of *τ*_*g*_(*b*) ∈ *S*^prim^. For each composite cell (*g, C*), Proposition D1 determines exactly one value of *α*_*g*_(*C*) ∈ *A*, Proposition D2 determines exactly one value of 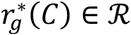, and Lemma E1 determines exactly one value of *m*_*g*_(*C*) ∈ ℳ.

Therefore

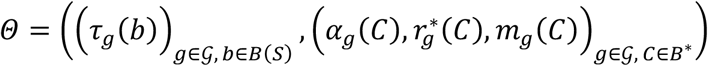

is one uniquely determined element of

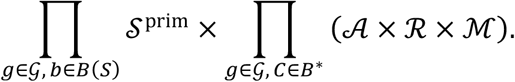

The fixed labelled topological inputs and conventions thus select one state for every locus-by-coordinate cell. This is fixed-input cellwise determinacy. It does not assert historical uniqueness or correctness of the supplied reference or empirical trees.

### E.3 Proof of Corollary 6

#### Proof of Corollary 6 (conditional numerical determinacy)

Assume that every normalized empirical tree carries a total nonnegative length function

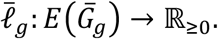

If τ_*g*_(*b*) = `Mapped`, the split 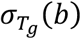 is displayed in 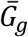 Lemma 2 gives one unique realizing empirical edge *e*_*b*_, so

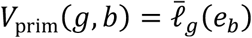

is single-valued. For every other primitive state, set *V*_prim_(*g, b*) = NA.

If 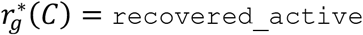, the active split is displayed and Lemma 2 gives one unique realizing edge *e*_*c*_. Hence

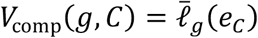

is single-valued.

If 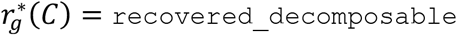, Lemma D2 supplies the unique finite component family *D*_*g*_(*C*). Every component split is displayed. Lemma D3 identifies one distinct component split for each component fiber, and Lemma 2 assigns each such split one unique empirical edge *e*_*F*_.

Therefore

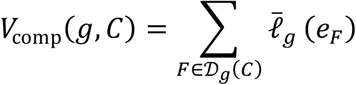

is a finite sum of uniquely selected values and is independent of enumeration order. For unrecovered composite cells, set *V*_comp_(*g, C*) = NA.

Collecting the primitive and composite values gives

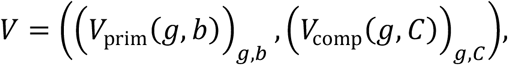

which is one uniquely determined element of

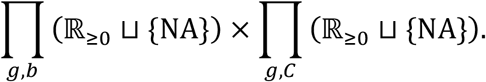

The index set is finite, but the value alphabet is not. Thus *V* is a finite-index numerical decoration whose determinacy is conditional on the supplied normalized lengths. The selected edges for a recovered decomposable coordinate need not be adjacent or form a continuous empirical path, and their sum is not asserted to be a full path length, a decomposition of a recovered active edge, or a sum of latent primitive-member lengths. No numerical value is claimed to be historical evolutionary truth.

## APPENDIX F. Boundary cases outside the main locus domain

### F.1 Two-taxon boundary proposition

#### Proposition F1 (two-taxon restriction and recovery)

Let *u, v* ∈ *T* be distinct and set

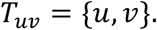

Let *P*_*S*_(*u, v*) be the unique path from *u* to *v* in *S* and define

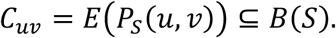

Then

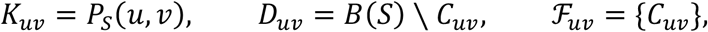

and

For every primitive edge *b*,

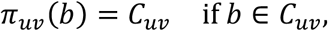

and

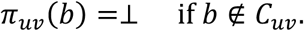

Moreover, |*C*_*uv*_| ≥ 2. Hence every primitive edge on the path is fused and has output state `NA_fuse`, every off-path edge is structural and has output state `NA_struct`, no primitive coordinate is eligible, and *C*_*uv*_ is the unique active composite coordinate of this restriction.

For empirical recovery, use boundary-local maps on the singleton domain {*C*_*uv*_} rather than the frozen global maps. Every normalized empirical tree on the leaf set {*u, v*} consists of exactly one edge and displays the split

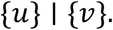

The active coordinate *C*_*uv*_ induces this same split and is therefore `recovered_active` with construction mode `single_edge_lookup`. If that empirical edge carries a valid normalized nonnegative length, the boundary-local numerical value is its length.

**Proof**. The minimal subtree connecting two retained leaves is their unique path, giving *K*_*uv*_ = *P*_0_(*u, v*) and *D*_*uv*_ = *B*(*S*) ∖ *C*_*uv*_. Every internal vertex of this path has degree two within *K*_*uv*_. Consecutive path edges are therefore adjacent in the degree-two edge-adjacency graph, so all edges of *C*_*uv*_ lie in one connected component. Since these are all edges of *K*_*uv*_, *F*_*uv*_ = {*C*_*uv*_} and the displayed form of π_*uv*_ follows.

If |*C*_*uv*_| = 1, the reference tree would contain one edge joining the two degree-one vertices *u* and *v*. Connectedness would then force *S* to consist only of that edge and to have two leaves, contradicting the standing assumption |*T*| ≥ 3. Thus |*C*_*uv*_| ≥ 2. The primitive-state conclusions follow immediately from Proposition 2.

A normalized unrooted tree on two labelled leaves has one edge and one split, {*u*} ∣ {*v*}. The unique active boundary coordinate induces that split, so it is recovered by one edge lookup. Numerical determinacy is conditional on a valid supplied length and does not decompose that value among the primitive path members.

The boundary coordinate does not alter the frozen cross-locus universe

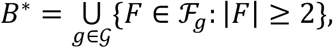

whose loci satisfy,*T*_*g*_, ≥ 3. Indeed, *C*_*uv*_ contains the terminal edges *b*_*u*_ and *b*_*v*_. If either taxon is absent from a main-domain locus, its terminal edge is deleted and *C*_*uv*_ cannot be a nondeleted fiber. If both are retained, their projected terminal splits are distinct because,*T*_*g*_, ≥ 3; Lemma 1 then prevents the two terminal edges from belonging to one fiber. Hence *C*_*uv*_ ∉ *B*^∗^ for the frozen locus family. The two-taxon construction uses boundary-local notation and does not enlarge *G*or *B*^∗^.

### F.2 One-taxon boundary remark

For *T*_*u*_ = {*u*}, the minimal connecting subtree is the single labelled vertex *u* and has no edges. Therefore

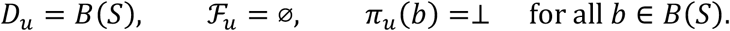

The retained-edge ledger and active-composite set are empty. The frozen primitive-axis row remains defined, and every primitive cell is `NA_struct`. There is no eligible coordinate and no empirical recovery step. The one-taxon case is a boundary remark and does not enter the main theorem domain.

