## Supplementary Computational Evidence for "A Unique Graph-Theoretic Truth Table for Cross-Gene Branch Identity": COMMANDS.md

# Replay commands

Run these commands from the extracted archive directory. They require Python 3.10 or later and use only the standard library.

```sh
mkdir replay

python3 scripts/run_and_log.py --log-dir replay/logs/static -- \
  python3 scripts/static_independence_check.py \
  scripts/run_validation_v0_2.py replay/audit/static

python3 scripts/run_and_log.py --log-dir replay/logs/run1 -- \
  python3 scripts/run_validation_v0_2.py --output-dir replay/run1

python3 scripts/run_and_log.py --log-dir replay/logs/run2 -- \
  python3 scripts/run_validation_v0_2.py --output-dir replay/run2

python3 scripts/run_and_log.py --log-dir replay/logs/qa -- \
  python3 scripts/qa_evidence_v0_2.py \
  --run-dir replay/run1 --rerun-dir replay/run2 --output-dir replay/audit/qa

shasum -a 256 -c SHA256SUMS.txt
```

Expected outcomes:

- both validation runs exit 0;
- the static audit reports 6/6 PASS;
- output-only QA reports 40/40 PASS;
- 13/13 canonical files compare byte-identically;
- `DISCREPANCIES.tsv` contains its header and zero data rows.
