## Supplementary Computational Evidence for "A Unique Graph-Theoretic Truth Table for Cross-Gene Branch Identity": INDEPENDENCE_AND_CIRCULARITY_AUDIT.md

# Independence and circularity audit v0.2

## Assessment

**PASS for the stated finite-domain implementation-fidelity claim.** The delivered graph- and split-side composite evaluators use separately implemented classification and recovery paths.

## Composite paths

Graph path:

1. Build all-pairs path union `K`.
2. Derive deleted original edges and degree-two fibers.
3. Classify composite state from deletion/fiber saturation.
4. Project graph-derived components to splits.
5. Query empirical splits and lengths through graph recovery logic.

Split path:

1. Project every original edge to the retained taxon set.
2. Mark empty projections structural and group equal nonempty projected splits.
3. Classify a composite through structural exclusion, whole/partial group intersections, and group coverage in `evaluate_split_composite()`.
4. Select component splits directly from the split-group index.
5. Query empirical splits and lengths within the split evaluator's own recovery branches.

The split evaluator does not call `graph_composite_state()`, `graph_composite_recovery()`, `graph_two_taxon_lookup()`, It does not receive graph composite results. Static check `S02` verifies this prohibited-call set.

## Boundary paths

The graph boundary evaluator derives a path fiber and graph-projected split. The split boundary evaluator reads the sole split group and performs its own lookup. The split boundary function calls neither the graph boundary nor either composite evaluator. Static check `S03` verifies this boundary-specific separation.

## Construction-order path

The fiber helper accepts an explicit edge set. In the order branch, its fourth argument is the locally computed `observed_k`; it no longer recomputes all-pairs restriction from the order's retained labels. The table exposes separate `k_edges_match` and `fibers_match` fields. Static checks `S04` and `S05` verify the call structure.

## Topology-count path

The set-partition recurrence is graph-free and does not call any topology generator. Output-only QA implements the same mathematical recurrence using different partition-enumeration code and does not import the validation script. Static check `S06` verifies absence of generator calls.

## Shared elements and residual circularity

The two sides share fixed coordinate labels, immutable sets/tuples, canonical split serialization/sorting, the same empirical topology, and the same declared length input. This sharing is necessary to compare answers on identical cases. Split groups are themselves the representation under comparison, so agreement after the group/fiber equivalence check is not probabilistic independence. The supported claim is narrower: separately implemented graph-side and split-side composite evaluators agree rowwise on the frozen domain.

AST checks cannot prove semantic correctness. Their role is to make prohibited function reuse machine-checkable; exhaustive rowwise comparison, output-only QA, and manual review supply the remaining evidence.
