## Supplementary Computational Evidence for "A Unique Graph-Theoretic Truth Table for Cross-Gene Branch Identity": MANUSCRIPT_RESULT_TABLE.md

# Proposed compact manuscript table

| Validation component | Enumerated scope | Result |
|---|---:|---:|
| Normalized empirical-tree domain | 472 cases: 260 binary, 212 multifurcating | Complete |
| Primitive graph/split comparison | 4,248 cells | 4,248 agreements |
| Separately implemented composite comparison | 9,912 cells | 9,912 agreements |
| Construction order using fibers derived from each observed `K` | 1,920 trials | 1,920 passed |
| Suppression order | 84 trials | 84 passed |
| Explicit two-leaf empirical lookup | 15 cases | 15 passed |
| Independent topology-count recurrence | 4 retained-taxon sizes | 1, 4, 26, 236; generator matched |
| Machine-readable discrepancies | All comparisons | 0 rows |
| Clean rerun / output-only QA / static audit | 13 files / 40 checks / 6 checks | 13 identical / 40 PASS / 6 PASS |
