## Supplementary Computational Evidence for "A Unique Graph-Theoretic Truth Table for Cross-Gene Branch Identity": QA_REPORT.md

# Independent QA report v0.2

## Decision

**PASS. Ready for independent review.**

The output-only QA script did not import or execute the validation implementation. It read the two persisted clean-run result trees, recomputed row counts and state totals, independently evaluated the set-partition recurrence, checked explicit boundary evidence, and compared hashes. All 40 checks passed.

The separate AST-based audit performed six code-path checks; all six passed.

## Reconciled evidence

- 472 unique main cases: 260 binary and 212 multifurcating.
- 4,248 unique primitive cells and 9,912 unique composite cells.
- 9,912/9,912 strengthened composite row agreements.
- 1,920/1,920 construction orders with both `K` and observed-`K-derived fibers matching.
- 84/84 suppression-order trials passing.
- 15/15 explicit two-leaf empirical lookups passing.
- Independent recurrence: `1, 4, 26, 236`, matching persisted recurrence and generated topologies.
- Zero discrepancy data rows; `DISCREPANCIES.tsv` contains its required header.
- 13/13 canonical files byte-identical across two fresh runs.
- The run-one output hash manifest matched all 12 files other than the manifest itself.

## State totals

All state totals reconcile to their row totals. Every claimed primitive, composite, recovery, topology-resolution, boundary, endpoint-collapse, and strengthened-check class has positive coverage.

## Independence qualification

Output-only QA verifies persisted evidence, not source semantics. The AST audit checks prohibited calls but is not a proof of algorithmic correctness. The independence conclusion therefore combines code inspection, static call-boundary checks, exhaustive rowwise agreement, and independent output reconciliation.

## Remaining limitations

This finite-domain QA does not prove the theorem, validate arbitrary reference trees, certify production software, or establish biological truth of the deterministic demonstration lengths.
