## Supplementary Computational Evidence for "A Unique Graph-Theoretic Truth Table for Cross-Gene Branch Identity": README.md

# Computational evidence for cross-gene branch identity

This archive accompanies *A Unique Graph-Theoretic Truth Table for Cross-Gene Branch Identity*. It contains the scripts, protocol, complete machine-readable outputs, discrepancy ledger, clean-rerun comparison, output-only QA, static call-boundary audit, and execution logs for the finite six-taxon computational validation reported in Section 7.

## Outcome

**PASS** with 0 discrepancy rows.

- 42 retained-taxon sets.
- 472 normalized empirical-tree cases: 260 binary and 212 multifurcating.
- 4,248 primitive graph/split comparisons.
- 9,912 separately implemented composite comparisons.
- 14,160 main categorical cells in total.
- 1,920 construction-order and 84 suppression-order trials.
- 15 explicit two-taxon empirical lookups and 6 one-taxon boundary cases.
- Two clean runs with byte-identical canonical outputs.
- 40/40 output-only QA checks passed.
- 6/6 static call-boundary checks passed.

## Quick replay

Requirements: Python 3.10 or later; standard library only.

From the extracted archive directory, follow `COMMANDS.md`. The replay writes only to a newly created `replay/` directory. Existing evidence files are not modified.

## Contents

- `scripts/`: validation, output-only QA, static call-boundary audit, and command logger.
- `outputs/`: canonical scientific outputs from the first clean run.
- `rerun/`: canonical outputs from the second clean run.
- `audit/`: output-only QA and static audit results.
- `logs/`: stdout, stderr, exact relative commands, exit status, and wall time.
- `VALIDATION_PROTOCOL_v0.2.md`: frozen validation design and pass/fail rules.
- `SCIENTIFIC_DESIGN.md`: compared constructions and failure policy.
- `INDEPENDENCE_AND_CIRCULARITY_AUDIT.md`: scope of evaluator separation.
- `QA_REPORT.md`: independent evidence reconciliation.
- `DISCREPANCIES.tsv`: required header and zero data rows.
- `SHA256SUMS.txt`: SHA-256 manifest for every other file in the extracted archive.

## Claim boundary

The computation checks implementation fidelity on a prespecified finite six-taxon domain. It does not prove the general theorem, certify production SplitAligner software, establish biological truth, or replace the mathematical proofs in the manuscript.
