## Supplementary Computational Evidence for "A Unique Graph-Theoretic Truth Table for Cross-Gene Branch Identity": SCIENTIFIC_DESIGN.md

# Scientific design v0.2

## Question

Do separately implemented graph- and split-side evaluators agree exhaustively on the prespecified finite six-tip domain?

## Unchanged domain

The fixed domain contains one reference tree, 42 retained sets, 472 normalized empirical topology cases, nine primitive coordinates, 21 composite member sets, 15 two-taxon cases, and 6 one-taxon cases. Multifurcating topologies remain included.

## Compared constructions

The graph path computes the unique minimal connecting subtree, deleted original edges, degree-two fibers, primitive states, graph composite states, and graph recovery.

The split path projects each original-edge split, groups equal nonempty projections, derives primitive states, and passes only split-derived structures into a separate composite evaluator. That evaluator classifies coordinates by structural/group coverage and queries the empirical split index directly for recovery and numerical construction.

Both sides intentionally share the coordinate universe, canonical split encoding, empirical topology, and declared length input. These are common problem inputs, not a reused composite classifier.

## Strengthened checks

- Composite comparison: 9,912 rowwise comparisons across reference state, saturation, components, component splits, recovery, mode, and value.
- Construction order: every observed ordered-anchor `K` is passed to the explicit-`K` fiber helper before comparison.
- Two-taxon recovery: each case uses an actual two-leaf empirical graph, split index, and length lookup.
- Topology counts: a graph-free set-partition recurrence supplies the expected counts independently of topology generation.

## Failure policy

Any row mismatch, missing state, failed recurrence, failed boundary lookup, failed static check, non-identical clean rerun forbids a success claim. The discrepancy table is generated from comparisons and retains a header even when it has zero data rows.
