## Supplementary Computational Evidence for "A Unique Graph-Theoretic Truth Table for Cross-Gene Branch Identity": qa_evidence_v0_2.py

```
#!/usr/bin/env python3
"""Independent, output-only QA for the branch-identity v0.2 evidence.

This script does not import or execute the validation implementation.  It
reconciles persisted TSV/JSON outputs, IDs, counts, states, discrepancies, and
clean-rerun byte hashes.
"""

from __future__ import annotations

import argparse
import csv
import hashlib
import json
from collections import Counter, defaultdict
from functools import lru_cache
from pathlib import Path
from typing import Iterable


def read_tsv(path: Path) -> list[dict[str, str]]:
    with path.open(encoding="utf-8", newline="") as handle:
        return list(csv.DictReader(handle, delimiter="\t"))


def sha256(path: Path) -> str:
    digest = hashlib.sha256()
    with path.open("rb") as handle:
        for chunk in iter(lambda: handle.read(1024 * 1024), b""):
            digest.update(chunk)
    return digest.hexdigest()


def relative_hashes(root: Path) -> dict[str, str]:
    return {
        str(path.relative_to(root)): sha256(path)
        for path in sorted(value for value in root.rglob("*") if value.is_file())
    }


def check(condition: bool, check_id: str, expected: object, observed: object, detail: str = "") -> dict[str, object]:
    return {
        "check_id": check_id,
        "status": "PASS" if condition else "FAIL",
        "expected": expected,
        "observed": observed,
        "detail": detail,
    }


def write_tsv(path: Path, rows: Iterable[dict[str, object]], fields: list[str]) -> None:
    path.parent.mkdir(parents=True, exist_ok=True)
    with path.open("w", encoding="utf-8", newline="") as handle:
        writer = csv.DictWriter(handle, fieldnames=fields, delimiter="\t", lineterminator="\n")
        writer.writeheader()
        for row in rows:
            writer.writerow({field: row.get(field, "") for field in fields})


def qa_set_partitions(values: tuple[int, ...]) -> Iterable[tuple[tuple[int, ...], ...]]:
    """Independent restricted-growth enumeration of unordered set partitions."""
    blocks: list[list[int]] = []

    def visit(index: int) -> Iterable[tuple[tuple[int, ...], ...]]:
        if index == len(values):
            yield tuple(tuple(block) for block in blocks)
            return
        value = values[index]
        for block in blocks:
            block.append(value)
            yield from visit(index + 1)
            block.pop()
        blocks.append([value])
        yield from visit(index + 1)
        blocks.pop()

    yield from visit(0)


@lru_cache(maxsize=None)
def qa_rooted_partition_count(values: tuple[int, ...]) -> int:
    if len(values) == 1:
        return 1
    total = 0
    for partition in qa_set_partitions(values):
        if len(partition) < 2:
            continue
        product = 1
        for block in partition:
            product *= qa_rooted_partition_count(tuple(block))
        total += product
    return total


def qa_unrooted_count(leaf_count: int) -> int:
    return qa_rooted_partition_count(tuple(range(leaf_count - 1)))


def main() -> int:
    parser = argparse.ArgumentParser()
    parser.add_argument("--run-dir", required=True)
    parser.add_argument("--rerun-dir", required=True)
    parser.add_argument("--output-dir", required=True)
    args = parser.parse_args()
    run = Path(args.run_dir)
    rerun = Path(args.rerun_dir)
    output = Path(args.output_dir)
    output.mkdir(parents=True, exist_ok=False)

    required = (
        "CASE_COUNT_RECONCILIATION.tsv",
        "STATE_COVERAGE.tsv",
        "INVARIANT_RESULTS.tsv",
        "DISCREPANCIES.tsv",
        "OUTPUT_SHA256SUMS.txt",
        "TOPOLOGY_COUNT_RECURRENCE.tsv",
        "outputs/MAIN_CASES.tsv",
        "outputs/RESTRICTION_RESULTS.tsv",
        "outputs/PRIMITIVE_RESULTS.tsv",
        "outputs/COMPOSITE_RESULTS.tsv",
        "outputs/ORDER_INVARIANCE.tsv",
        "outputs/BOUNDARY_RESULTS.tsv",
        "outputs/SUMMARY.json",
    )
    checks: list[dict[str, object]] = []
    missing = [value for value in required if not (run / value).is_file()]
    checks.append(check(not missing, "Q01_required_files", 0, len(missing), ",".join(missing)))
    if missing:
        (output / "QA_AUDIT.json").write_text(json.dumps({"status": "FAIL", "checks": checks}, indent=2) + "\n")
        return 1

    cases = read_tsv(run / "outputs/MAIN_CASES.tsv")
    restrictions = read_tsv(run / "outputs/RESTRICTION_RESULTS.tsv")
    primitive = read_tsv(run / "outputs/PRIMITIVE_RESULTS.tsv")
    composite = read_tsv(run / "outputs/COMPOSITE_RESULTS.tsv")
    order = read_tsv(run / "outputs/ORDER_INVARIANCE.tsv")
    boundary = read_tsv(run / "outputs/BOUNDARY_RESULTS.tsv")
    counts = read_tsv(run / "CASE_COUNT_RECONCILIATION.tsv")
    coverage = read_tsv(run / "STATE_COVERAGE.tsv")
    invariants = read_tsv(run / "INVARIANT_RESULTS.tsv")
    discrepancies = read_tsv(run / "DISCREPANCIES.tsv")
    summary = json.loads((run / "outputs/SUMMARY.json").read_text(encoding="utf-8"))
    recurrence = read_tsv(run / "TOPOLOGY_COUNT_RECURRENCE.tsv")

    checks.extend(
        [
            check(len(cases) == 472, "Q02_main_cases", 472, len(cases)),
            check(len({row["case_id"] for row in cases}) == 472, "Q03_unique_case_ids", 472, len({row["case_id"] for row in cases})),
            check(len(restrictions) == 42, "Q04_restriction_rows", 42, len(restrictions)),
            check(len({row["retained_id"] for row in restrictions}) == 42, "Q05_unique_retained_ids", 42, len({row["retained_id"] for row in restrictions})),
            check(len(primitive) == 4248, "Q06_primitive_rows", 4248, len(primitive)),
            check(len({row["cell_id"] for row in primitive}) == 4248, "Q07_unique_primitive_ids", 4248, len({row["cell_id"] for row in primitive})),
            check(len(composite) == 9912, "Q08_composite_rows", 9912, len(composite)),
            check(len({row["cell_id"] for row in composite}) == 9912, "Q09_unique_composite_ids", 9912, len({row["cell_id"] for row in composite})),
            check(not ({row["cell_id"] for row in primitive} & {row["cell_id"] for row in composite}), "Q10_cross_type_id_disjoint", 0, len({row["cell_id"] for row in primitive} & {row["cell_id"] for row in composite})),
        ]
    )

    cases_by_retained = Counter(row["retained_count"] for row in cases)
    checks.extend(
        [
            check(cases_by_retained == Counter({"3": 20, "4": 60, "5": 156, "6": 236}), "Q11_case_count_by_retained_size", {"3": 20, "4": 60, "5": 156, "6": 236}, dict(cases_by_retained)),
            check(Counter(row["resolution_class"] for row in cases) == Counter({"binary": 260, "multifurcating": 212}), "Q12_resolution_counts", {"binary": 260, "multifurcating": 212}, dict(Counter(row["resolution_class"] for row in cases))),
            check(all(value == 9 for value in Counter(row["case_id"] for row in primitive).values()), "Q13_nine_primitive_rows_per_case", "all 9", sorted(set(Counter(row["case_id"] for row in primitive).values()))),
            check(all(value == 21 for value in Counter(row["case_id"] for row in composite).values()), "Q14_twentyone_composite_rows_per_case", "all 21", sorted(set(Counter(row["case_id"] for row in composite).values()))),
        ]
    )

    primitive_agreement = sum(
        row["graph_reference_state"] == row["split_reference_state"]
        and row["graph_fiber"] == row["split_group"]
        and row["graph_output_state"] == row["split_output_state"]
        for row in primitive
    )
    composite_agreement = sum(
        row["graph_reference_state"] == row["split_reference_state"]
        and row["graph_saturated"] == row["split_saturated"]
        and row["graph_components"] == row["split_components"]
        and row["graph_component_splits"] == row["split_component_splits"]
        and row["graph_recovery_state"] == row["split_recovery_state"]
        and row["graph_construction_mode"] == row["split_construction_mode"]
        and row["graph_numeric_value"] == row["split_numeric_value"]
        for row in composite
    )
    checks.extend(
        [
            check(primitive_agreement == 4248, "Q15_primitive_oracle_comparator_agreement", 4248, primitive_agreement),
            check(composite_agreement == 9912, "Q16_composite_oracle_comparator_agreement", 9912, composite_agreement),
            check(sum(row["pass"] == "true" for row in order) == 2004, "Q17_order_trials_pass", 2004, sum(row["pass"] == "true" for row in order)),
            check(Counter(row["trial_type"] for row in order) == Counter({"construction_order": 1920, "suppression_order": 84}), "Q18_order_trial_counts", {"construction_order": 1920, "suppression_order": 84}, dict(Counter(row["trial_type"] for row in order))),
        ]
    )

    boundary_primitive = [row for row in boundary if row["row_type"] == "primitive"]
    boundary_composite = [row for row in boundary if row["row_type"] == "composite"]
    checks.extend(
        [
            check(len(boundary_primitive) == 189, "Q19_boundary_primitive_rows", 189, len(boundary_primitive)),
            check(len(boundary_composite) == 15, "Q20_boundary_composite_rows", 15, len(boundary_composite)),
            check(all(row["pass"] == "true" for row in boundary), "Q21_boundary_all_pass", "all true", sum(row["pass"] != "true" for row in boundary)),
            check(all(row["graph_recovery_state"] == row["split_recovery_state"] == "recovered_active" and row["graph_construction_mode"] == row["split_construction_mode"] == "single_edge_lookup" and row["graph_numeric_value"] == row["split_numeric_value"] == "1" for row in boundary_composite), "Q22_two_taxon_composite_recovery", "all independent lookups recovered_active/single_edge_lookup/1", sum(not (row["graph_recovery_state"] == row["split_recovery_state"] == "recovered_active" and row["graph_construction_mode"] == row["split_construction_mode"] == "single_edge_lookup" and row["graph_numeric_value"] == row["split_numeric_value"] == "1") for row in boundary_composite)),
        ]
    )

    checks.extend(
        [
            check(not discrepancies, "Q23_discrepancy_rows", 0, len(discrepancies)),
            check(all(row["pass"] == "true" and row["expected"] == row["observed"] for row in counts), "Q24_count_reconciliation", "all pass/equal", sum(not (row["pass"] == "true" and row["expected"] == row["observed"]) for row in counts)),
            check(all(row["pass"] == "true" and int(row["observed_count"]) >= int(row["required_minimum"]) for row in coverage), "Q25_state_coverage", "all pass", sum(row["pass"] != "true" for row in coverage)),
            check(all(row["pass"] == "true" for row in invariants), "Q26_invariants", "all pass", sum(row["pass"] != "true" for row in invariants)),
            check(summary.get("status") == "PASS" and summary.get("discrepancy_count") == 0, "Q27_summary_status", "PASS/0", f"{summary.get('status')}/{summary.get('discrepancy_count')}")
        ]
    )

    primitive_state_counts = Counter(row["graph_output_state"] for row in primitive)
    composite_reference_counts = Counter(row["graph_reference_state"] for row in composite)
    composite_recovery_counts = Counter(row["graph_recovery_state"] for row in composite)
    checks.extend(
        [
            check(sum(primitive_state_counts.values()) == 4248, "Q28_primitive_state_sum", 4248, sum(primitive_state_counts.values()), json.dumps(dict(primitive_state_counts), sort_keys=True)),
            check(sum(composite_reference_counts.values()) == 9912, "Q29_composite_reference_sum", 9912, sum(composite_reference_counts.values()), json.dumps(dict(composite_reference_counts), sort_keys=True)),
            check(sum(composite_recovery_counts.values()) == 9912, "Q30_composite_recovery_sum", 9912, sum(composite_recovery_counts.values()), json.dumps(dict(composite_recovery_counts), sort_keys=True)),
        ]
    )

    run_hashes = relative_hashes(run)
    rerun_hashes = relative_hashes(rerun)
    all_paths = sorted(set(run_hashes) | set(rerun_hashes))
    comparison_rows = []
    for relative in all_paths:
        first = run_hashes.get(relative, "MISSING")
        second = rerun_hashes.get(relative, "MISSING")
        comparison_rows.append(
            {
                "relative_path": relative,
                "run1_sha256": first,
                "run2_sha256": second,
                "byte_identical": str(first == second).lower(),
            }
        )
    identical_count = sum(row["byte_identical"] == "true" for row in comparison_rows)
    checks.append(check(identical_count == len(comparison_rows), "Q31_clean_rerun_byte_identity", len(comparison_rows), identical_count))

    declared_hashes = {}
    for line in (run / "OUTPUT_SHA256SUMS.txt").read_text(encoding="utf-8").splitlines():
        digest, relative = line.split("  ", 1)
        declared_hashes[relative] = digest
    actual_without_manifest = {
        relative: digest for relative, digest in run_hashes.items() if relative != "OUTPUT_SHA256SUMS.txt"
    }
    checks.append(check(declared_hashes == actual_without_manifest, "Q32_output_hash_manifest", len(actual_without_manifest), len(declared_hashes)))

    independent_recurrence = {size: qa_unrooted_count(size) for size in range(3, 7)}
    recurrence_by_size = {int(row["retained_taxa_count"]): row for row in recurrence}
    construction_rows = [row for row in order if row["trial_type"] == "construction_order"]
    strengthened_coverage = {
        row["state_or_class"]: int(row["observed_count"])
        for row in coverage
        if row["coverage_family"] == "strengthened_check"
    }
    expected_strengthened = {
        "independent_split_composite_rows": 9912,
        "explicit_two_leaf_empirical_lookups": 15,
        "observed_k_fiber_trials": 1920,
        "partition_recurrence_sizes": 4,
    }
    summary_strengthened = summary.get("strengthened_checks", {})
    recurrence_count_rows = {
        row["count_id"]: row for row in counts if row["count_id"].startswith("topology_recurrence_k")
    }
    checks.extend(
        [
            check(len(recurrence) == 4 and set(recurrence_by_size) == {3, 4, 5, 6}, "Q33_recurrence_rows", "four sizes 3-6", sorted(recurrence_by_size)),
            check(all(int(recurrence_by_size[size]["partition_recurrence_count"]) == independent_recurrence[size] for size in range(3, 7)), "Q34_independent_recurrence_matches_table", independent_recurrence, {size: int(recurrence_by_size[size]["partition_recurrence_count"]) for size in recurrence_by_size}),
            check(independent_recurrence == {3: 1, 4: 4, 5: 26, 6: 236}, "Q35_independent_recurrence_sequence", {3: 1, 4: 4, 5: 26, 6: 236}, independent_recurrence),
            check(len(construction_rows) == 1920 and all(row["k_edges_match"] == "true" for row in construction_rows), "Q36_construction_observed_k_match", 1920, sum(row["k_edges_match"] == "true" for row in construction_rows)),
            check(len(construction_rows) == 1920 and all(row["fibers_match"] == "true" for row in construction_rows), "Q37_construction_observed_k_fibers_match", 1920, sum(row["fibers_match"] == "true" for row in construction_rows)),
            check(all(row["empirical_edge_count"] == "1" and row["graph_lookup_split"] == row["split_lookup_split"] == row["empirical_topology_signature"] and row["declared_length_rule"] == 'single empirical edge assigned Decimal("1")' for row in boundary_composite), "Q38_explicit_two_leaf_empirical_evidence", "15 explicit one-edge split lookups", sum(row["empirical_edge_count"] == "1" and row["graph_lookup_split"] == row["split_lookup_split"] == row["empirical_topology_signature"] and row["declared_length_rule"] == 'single empirical edge assigned Decimal("1")' for row in boundary_composite)),
            check(strengthened_coverage == expected_strengthened, "Q39_strengthened_coverage", expected_strengthened, strengthened_coverage),
            check(len(recurrence_count_rows) == 4 and all(row["pass"] == "true" and int(row["observed"]) == independent_recurrence[int(row["count_id"].removeprefix("topology_recurrence_k"))] for row in recurrence_count_rows.values()) and summary_strengthened == {"explicit_two_leaf_empirical_lookups": 15, "independent_split_composite_rows": 9912, "observed_k_fiber_trials": 1920, "topology_partition_recurrence_sizes": 4}, "Q40_strengthened_summary_and_count_rows", "four recurrence rows and strengthened summary", {"recurrence_rows": len(recurrence_count_rows), "summary": summary_strengthened}),
        ]
    )

    failed = [row for row in checks if row["status"] != "PASS"]
    audit = {
        "status": "PASS" if not failed else "FAIL",
        "check_count": len(checks),
        "failed_check_count": len(failed),
        "checks": checks,
        "independence_note": "Output-only QA; validation implementation was not imported.",
        "observed_state_counts": {
            "primitive_output": dict(sorted(primitive_state_counts.items())),
            "composite_reference": dict(sorted(composite_reference_counts.items())),
            "composite_recovery": dict(sorted(composite_recovery_counts.items())),
        },
    }
    (output / "QA_AUDIT.json").write_text(json.dumps(audit, indent=2, sort_keys=True) + "\n", encoding="utf-8")
    write_tsv(
        output / "QA_CHECKS.tsv",
        checks,
        ["check_id", "status", "expected", "observed", "detail"],
    )
    write_tsv(
        output / "RERUN_HASH_COMPARISON.tsv",
        comparison_rows,
        ["relative_path", "run1_sha256", "run2_sha256", "byte_identical"],
    )
    print(json.dumps({"status": audit["status"], "check_count": len(checks), "failed": len(failed), "rerun_files": len(comparison_rows)}, sort_keys=True))
    return 0 if not failed else 1


if __name__ == "__main__":
    raise SystemExit(main())
```
