## Supplementary Computational Evidence for "A Unique Graph-Theoretic Truth Table for Cross-Gene Branch Identity": run_and_log.py

```
#!/usr/bin/env python3
"""Run one command and persist exact command, streams, exit code, and wall time."""

from __future__ import annotations

import argparse
import json
import shlex
import subprocess
import time
from pathlib import Path


def main() -> int:
    parser = argparse.ArgumentParser()
    parser.add_argument("--log-dir", required=True)
    parser.add_argument("command", nargs=argparse.REMAINDER)
    args = parser.parse_args()
    command = args.command
    if command and command[0] == "--":
        command = command[1:]
    if not command:
        raise SystemExit("missing command")
    log_dir = Path(args.log_dir)
    log_dir.mkdir(parents=True, exist_ok=False)
    start = time.perf_counter()
    completed = subprocess.run(command, text=True, capture_output=True, check=False)
    wall = time.perf_counter() - start
    (log_dir / "stdout.log").write_text(completed.stdout, encoding="utf-8")
    (log_dir / "stderr.log").write_text(completed.stderr, encoding="utf-8")
    record = {
        "command_argv": command,
        "command_shell_escaped": shlex.join(command),
        "exit_code": completed.returncode,
        "wall_time_seconds": round(wall, 6),
    }
    (log_dir / "run_record.json").write_text(
        json.dumps(record, indent=2, sort_keys=True) + "\n", encoding="utf-8"
    )
    print(json.dumps(record, sort_keys=True))
    return completed.returncode


if __name__ == "__main__":
    raise SystemExit(main())
```
