## Supplementary Computational Evidence for "A Unique Graph-Theoretic Truth Table for Cross-Gene Branch Identity": run_validation_v0_2.py

```
#!/usr/bin/env python3
"""Deterministic exhaustive validation for branch-identity validation protocol v0.2.

The graph oracle builds restriction fibers only from paths, vertex degrees, and
edge-component connectivity.  The split comparator independently projects
reference-edge splits, groups equal projections, evaluates composite states and
recovery through split-index queries, and never calls graph composite logic.
"""

from __future__ import annotations

import argparse
import csv
import hashlib
import itertools
import json
from collections import Counter, defaultdict, deque
from dataclasses import dataclass
from decimal import Decimal
from functools import lru_cache
from pathlib import Path
from typing import Iterable, Mapping, Sequence


TAXA = ("A", "B", "C", "D", "E", "F")
COORDS = ("A", "B", "C", "D", "E", "F", "N_BC", "N_DEF", "N_EF")
COORD_INDEX = {value: index for index, value in enumerate(COORDS)}
EXPECTED_TOPOLOGY_COUNTS = {3: 1, 4: 4, 5: 26, 6: 236}
EXPECTED_BINARY_COUNTS = {3: 1, 4: 3, 5: 15, 6: 105}
EXPECTED_COMPOSITE_AXIS = {
    frozenset(value.split("|"))
    for value in (
        "A|N_BC",
        "A|N_DEF",
        "B|N_BC",
        "C|N_BC",
        "D|N_DEF",
        "D|N_EF",
        "E|N_EF",
        "F|N_EF",
        "N_BC|N_DEF",
        "N_DEF|N_EF",
        "A|N_DEF|N_EF",
        "B|N_BC|N_DEF",
        "C|N_BC|N_DEF",
        "D|N_BC|N_DEF",
        "E|N_DEF|N_EF",
        "F|N_DEF|N_EF",
        "N_BC|N_DEF|N_EF",
        "B|N_BC|N_DEF|N_EF",
        "C|N_BC|N_DEF|N_EF",
        "E|N_BC|N_DEF|N_EF",
        "F|N_BC|N_DEF|N_EF",
    )
}


def edge_key(a: str, b: str) -> tuple[str, str]:
    return (a, b) if a < b else (b, a)


def coord_sort(values: Iterable[str]) -> tuple[int, ...]:
    return tuple(COORD_INDEX[value] for value in sorted(values, key=COORD_INDEX.__getitem__))


def member_key(values: Iterable[str]) -> str:
    return "|".join(sorted(set(values), key=COORD_INDEX.__getitem__))


def member_sets_text(values: Iterable[frozenset[str]]) -> str:
    return ";".join(member_key(value) for value in sorted(values, key=coord_sort))


def taxa_key(values: Iterable[str]) -> str:
    return "".join(sorted(values))


def canonical_split(left: Iterable[str], right: Iterable[str]) -> tuple[tuple[str, ...], tuple[str, ...]]:
    a = tuple(sorted(set(left)))
    b = tuple(sorted(set(right)))
    if not a or not b:
        raise ValueError("split sides must be nonempty")
    return (a, b) if a <= b else (b, a)


def split_key(split: tuple[tuple[str, ...], tuple[str, ...]]) -> str:
    return ",".join(split[0]) + "||" + ",".join(split[1])


class Graph:
    def __init__(self) -> None:
        self.adj: dict[str, set[str]] = defaultdict(set)
        self.labels: dict[tuple[str, str], str] = {}

    def add_edge(self, a: str, b: str, label: str = "") -> None:
        edge = edge_key(a, b)
        if b in self.adj[a]:
            raise ValueError(f"duplicate edge {edge}")
        self.adj[a].add(b)
        self.adj[b].add(a)
        if label:
            self.labels[edge] = label

    def remove_edge(self, a: str, b: str) -> None:
        edge = edge_key(a, b)
        self.adj[a].remove(b)
        self.adj[b].remove(a)
        self.labels.pop(edge, None)

    @property
    def nodes(self) -> set[str]:
        return set(self.adj)

    @property
    def edges(self) -> set[tuple[str, str]]:
        return {edge_key(a, b) for a, neighbours in self.adj.items() for b in neighbours if a < b}

    def copy(self) -> "Graph":
        result = Graph()
        for edge in sorted(self.edges):
            result.add_edge(edge[0], edge[1], self.labels.get(edge, ""))
        return result


def reference_graph() -> tuple[Graph, dict[str, tuple[str, str]]]:
    graph = Graph()
    definitions = (
        ("X", "A", "A"),
        ("Y", "B", "B"),
        ("Y", "C", "C"),
        ("Z", "D", "D"),
        ("W", "E", "E"),
        ("W", "F", "F"),
        ("X", "Y", "N_BC"),
        ("X", "Z", "N_DEF"),
        ("Z", "W", "N_EF"),
    )
    coordinate_to_edge: dict[str, tuple[str, str]] = {}
    for a, b, coordinate in definitions:
        graph.add_edge(a, b, coordinate)
        coordinate_to_edge[coordinate] = edge_key(a, b)
    return graph, coordinate_to_edge


def unique_path_edges(graph: Graph, start: str, end: str) -> list[tuple[str, str]]:
    parent: dict[str, str | None] = {start: None}
    queue: deque[str] = deque([start])
    while queue:
        node = queue.popleft()
        if node == end:
            break
        for nxt in sorted(graph.adj[node]):
            if nxt not in parent:
                parent[nxt] = node
                queue.append(nxt)
    if end not in parent:
        raise ValueError("disconnected graph")
    edges: list[tuple[str, str]] = []
    node = end
    while parent[node] is not None:
        previous = parent[node]
        assert previous is not None
        edges.append(edge_key(node, previous))
        node = previous
    edges.reverse()
    return edges


def component_taxa_without_edge(
    graph: Graph, edge: tuple[str, str], leaf_set: set[str]
) -> tuple[set[str], set[str]]:
    start = edge[0]
    seen = {start}
    stack = [start]
    while stack:
        node = stack.pop()
        for nxt in graph.adj[node]:
            if edge_key(node, nxt) == edge or nxt in seen:
                continue
            seen.add(nxt)
            stack.append(nxt)
    left = seen & leaf_set
    return left, leaf_set - left


def projected_split(
    graph: Graph, edge: tuple[str, str], retained: set[str]
) -> tuple[tuple[str, ...], tuple[str, ...]] | None:
    left, right = component_taxa_without_edge(graph, edge, retained)
    if not left or not right:
        return None
    return canonical_split(left, right)


def empirical_split_index(graph: Graph, leaves: set[str]) -> dict[tuple[tuple[str, ...], tuple[str, ...]], tuple[str, str]]:
    result: dict[tuple[tuple[str, ...], tuple[str, ...]], tuple[str, str]] = {}
    for edge in sorted(graph.edges):
        split = projected_split(graph, edge, leaves)
        if split is None:
            raise AssertionError("empirical edge produced an empty split side")
        if split in result:
            raise AssertionError("empirical edge-split map is not injective")
        result[split] = edge
    return result


def connected_edge_components(
    edges: set[tuple[str, str]], adjacency: Mapping[tuple[str, str], set[tuple[str, str]]]
) -> list[set[tuple[str, str]]]:
    unseen = set(edges)
    result: list[set[tuple[str, str]]] = []
    while unseen:
        seed = min(unseen)
        unseen.remove(seed)
        component = {seed}
        stack = [seed]
        while stack:
            edge = stack.pop()
            for nxt in sorted(adjacency[edge]):
                if nxt in unseen:
                    unseen.remove(nxt)
                    component.add(nxt)
                    stack.append(nxt)
        result.append(component)
    return result


@dataclass(frozen=True)
class Restriction:
    retained: frozenset[str]
    k_edges: frozenset[tuple[str, str]]
    deleted: frozenset[str]
    fibers: tuple[frozenset[str], ...]
    coordinate_to_fiber: Mapping[str, frozenset[str] | None]
    reference_state: Mapping[str, str]


def restriction_from_k_edges(
    reference: Graph,
    coordinate_to_edge: Mapping[str, tuple[str, str]],
    retained_values: Iterable[str],
    supplied_k_edges: Iterable[tuple[str, str]],
) -> Restriction:
    """Derive deletion, fibers, and states from the exact supplied K edge set."""
    retained = frozenset(retained_values)
    k_edges = {edge_key(*edge) for edge in supplied_k_edges}
    unknown_edges = k_edges - reference.edges
    if unknown_edges:
        raise ValueError(f"supplied K contains non-reference edges: {sorted(unknown_edges)}")
    degrees: Counter[str] = Counter()
    incident: dict[str, list[tuple[str, str]]] = defaultdict(list)
    for edge in k_edges:
        for node in edge:
            degrees[node] += 1
            incident[node].append(edge)
    edge_adjacency: dict[tuple[str, str], set[tuple[str, str]]] = {edge: set() for edge in k_edges}
    for node, node_edges in incident.items():
        if degrees[node] == 2:
            first, second = node_edges
            edge_adjacency[first].add(second)
            edge_adjacency[second].add(first)
    edge_to_coordinate = {edge: coordinate for coordinate, edge in coordinate_to_edge.items()}
    fiber_values = [
        frozenset(edge_to_coordinate[edge] for edge in component)
        for component in connected_edge_components(k_edges, edge_adjacency)
    ]
    fibers = tuple(sorted(fiber_values, key=coord_sort))
    deleted = frozenset(
        coordinate for coordinate, edge in coordinate_to_edge.items() if edge not in k_edges
    )
    coordinate_to_fiber: dict[str, frozenset[str] | None] = {coordinate: None for coordinate in COORDS}
    for fiber in fibers:
        for coordinate in fiber:
            coordinate_to_fiber[coordinate] = fiber
    states: dict[str, str] = {}
    for coordinate in COORDS:
        fiber = coordinate_to_fiber[coordinate]
        if fiber is None:
            states[coordinate] = "structural"
        elif len(fiber) >= 2:
            states[coordinate] = "fused"
        else:
            states[coordinate] = "eligible"
    return Restriction(retained, frozenset(k_edges), deleted, fibers, coordinate_to_fiber, states)


def graph_restriction(
    reference: Graph, coordinate_to_edge: Mapping[str, tuple[str, str]], retained_values: Iterable[str]
) -> Restriction:
    retained = frozenset(retained_values)
    k_edges: set[tuple[str, str]] = set()
    for a, b in itertools.combinations(sorted(retained), 2):
        k_edges.update(unique_path_edges(reference, a, b))
    return restriction_from_k_edges(reference, coordinate_to_edge, retained, k_edges)


@dataclass(frozen=True)
class SplitRestriction:
    structural: frozenset[str]
    groups: tuple[frozenset[str], ...]
    coordinate_to_group: Mapping[str, frozenset[str] | None]
    group_to_split: Mapping[frozenset[str], tuple[tuple[str, ...], tuple[str, ...]]]
    reference_state: Mapping[str, str]


def split_restriction(
    reference: Graph, coordinate_to_edge: Mapping[str, tuple[str, str]], retained_values: Iterable[str]
) -> SplitRestriction:
    retained = set(retained_values)
    grouped: dict[tuple[tuple[str, ...], tuple[str, ...]], set[str]] = defaultdict(set)
    structural: set[str] = set()
    for coordinate in COORDS:
        split = projected_split(reference, coordinate_to_edge[coordinate], retained)
        if split is None:
            structural.add(coordinate)
        else:
            grouped[split].add(coordinate)
    groups = tuple(sorted((frozenset(values) for values in grouped.values()), key=coord_sort))
    coordinate_to_group: dict[str, frozenset[str] | None] = {coordinate: None for coordinate in COORDS}
    group_to_split: dict[frozenset[str], tuple[tuple[str, ...], tuple[str, ...]]] = {}
    for split, values in grouped.items():
        group = frozenset(values)
        group_to_split[group] = split
        for coordinate in group:
            coordinate_to_group[coordinate] = group
    states = {}
    for coordinate in COORDS:
        group = coordinate_to_group[coordinate]
        if group is None:
            states[coordinate] = "structural"
        elif len(group) >= 2:
            states[coordinate] = "fused"
        else:
            states[coordinate] = "eligible"
    return SplitRestriction(
        frozenset(structural), groups, coordinate_to_group, group_to_split, states
    )


def k_edges_from_order(reference: Graph, order: Sequence[str]) -> set[tuple[str, str]]:
    if len(order) < 2:
        return set()
    anchor = order[0]
    result: set[tuple[str, str]] = set()
    for taxon in order[1:]:
        result.update(unique_path_edges(reference, anchor, taxon))
    return result


def suppression_partition(
    reference: Graph, coordinate_to_edge: Mapping[str, tuple[str, str]], k_edges: set[tuple[str, str]], reverse: bool
) -> tuple[frozenset[str], ...]:
    adjacency: dict[str, set[str]] = defaultdict(set)
    provenance: dict[tuple[str, str], frozenset[str]] = {}
    edge_to_coordinate = {edge: coordinate for coordinate, edge in coordinate_to_edge.items()}
    for edge in k_edges:
        adjacency[edge[0]].add(edge[1])
        adjacency[edge[1]].add(edge[0])
        provenance[edge] = frozenset({edge_to_coordinate[edge]})
    while True:
        candidates = sorted((node for node, neighbours in adjacency.items() if len(neighbours) == 2), reverse=reverse)
        if not candidates:
            break
        node = candidates[0]
        a, b = sorted(adjacency[node])
        first = edge_key(a, node)
        second = edge_key(b, node)
        combined = provenance.pop(first) | provenance.pop(second)
        adjacency[a].remove(node)
        adjacency[b].remove(node)
        del adjacency[node]
        merged = edge_key(a, b)
        if merged in provenance:
            raise AssertionError("suppression created a duplicate tree edge")
        adjacency[a].add(b)
        adjacency[b].add(a)
        provenance[merged] = combined
    return tuple(sorted(provenance.values(), key=coord_sort))


def labelled_set_partitions(values: tuple[str, ...]) -> Iterable[tuple[tuple[str, ...], ...]]:
    """Enumerate each unordered set partition once in canonical block order."""
    if not values:
        yield ()
        return
    first = values[0]
    for partition in labelled_set_partitions(values[1:]):
        yield ((first,),) + partition
        for index, block in enumerate(partition):
            yield partition[:index] + ((first,) + block,) + partition[index + 1 :]


@lru_cache(maxsize=None)
def rooted_partition_tree_count(labels: tuple[str, ...]) -> int:
    """Count rooted nonbinary phylogenetic trees by the root-child partition recurrence."""
    if len(labels) == 1:
        return 1
    total = 0
    for partition in labelled_set_partitions(labels):
        if len(partition) < 2:
            continue
        product = 1
        for block in partition:
            product *= rooted_partition_tree_count(tuple(block))
        total += product
    return total


def unrooted_topology_count_recurrence(leaf_count: int) -> int:
    """Use U(n)=R(n-1), after removing a distinguished leaf and rooting at its neighbour."""
    if leaf_count < 3:
        raise ValueError("unrooted normalized topology recurrence requires at least three leaves")
    rooted_labels = tuple(f"L{index}" for index in range(1, leaf_count))
    return rooted_partition_tree_count(rooted_labels)


def topology_signature(graph: Graph, leaves: set[str]) -> tuple[str, ...]:
    return tuple(sorted(split_key(split) for split in empirical_split_index(graph, leaves)))


def insert_leaf(graph: Graph, edge: tuple[str, str], leaf: str, serial: int) -> Graph:
    result = graph.copy()
    result.remove_edge(*edge)
    internal = f"I_{leaf}_{serial}"
    while internal in result.nodes:
        internal += "x"
    result.add_edge(edge[0], internal)
    result.add_edge(edge[1], internal)
    result.add_edge(internal, leaf)
    return result


def binary_topologies(labels: tuple[str, ...]) -> dict[tuple[str, ...], Graph]:
    if len(labels) < 3:
        raise ValueError("binary topology generator requires at least three leaves")
    base = Graph()
    base.add_edge("I0", labels[0])
    base.add_edge("I0", labels[1])
    base.add_edge("I0", labels[2])
    current = {topology_signature(base, set(labels[:3])): base}
    for leaf in labels[3:]:
        next_values: dict[tuple[str, ...], Graph] = {}
        serial = 0
        for graph in (current[key] for key in sorted(current)):
            for edge in sorted(graph.edges):
                serial += 1
                candidate = insert_leaf(graph, edge, leaf, serial)
                signature = topology_signature(candidate, set(labels[: labels.index(leaf) + 1]))
                next_values.setdefault(signature, candidate)
        current = next_values
    return current


class UnionFind:
    def __init__(self, values: Iterable[str]) -> None:
        self.parent = {value: value for value in values}

    def find(self, value: str) -> str:
        while self.parent[value] != value:
            self.parent[value] = self.parent[self.parent[value]]
            value = self.parent[value]
        return value

    def union(self, a: str, b: str) -> None:
        ra, rb = self.find(a), self.find(b)
        if ra != rb:
            self.parent[max(ra, rb)] = min(ra, rb)


def contract_internal_subset(graph: Graph, leaves: set[str], selected: Sequence[tuple[str, str]]) -> Graph:
    union = UnionFind(graph.nodes)
    for edge in selected:
        union.union(*edge)
    classes: dict[str, set[str]] = defaultdict(set)
    for node in graph.nodes:
        classes[union.find(node)].add(node)
    representative: dict[str, str] = {}
    for root, values in classes.items():
        leaf_values = values & leaves
        if leaf_values:
            if len(leaf_values) != 1 or len(values) != 1:
                raise AssertionError("internal-edge contraction merged a leaf")
            name = next(iter(leaf_values))
        else:
            name = "Q[" + "+".join(sorted(values)) + "]"
        for value in values:
            representative[value] = name
    result = Graph()
    for edge in sorted(graph.edges):
        a, b = representative[edge[0]], representative[edge[1]]
        if a != b and b not in result.adj[a]:
            result.add_edge(a, b)
    return result


def all_normalized_topologies(labels: tuple[str, ...]) -> dict[tuple[str, ...], Graph]:
    leaves = set(labels)
    result: dict[tuple[str, ...], Graph] = {}
    binaries = binary_topologies(labels)
    for signature in sorted(binaries):
        binary = binaries[signature]
        internal_edges = sorted(edge for edge in binary.edges if edge[0] not in leaves and edge[1] not in leaves)
        for mask in range(1 << len(internal_edges)):
            selected = [edge for index, edge in enumerate(internal_edges) if mask & (1 << index)]
            candidate = contract_internal_subset(binary, leaves, selected)
            for node in candidate.nodes - leaves:
                if len(candidate.adj[node]) < 3:
                    raise AssertionError("normalized topology has internal degree below three")
            candidate_signature = topology_signature(candidate, leaves)
            result.setdefault(candidate_signature, candidate)
    return result


def topology_is_binary(graph: Graph, leaves: set[str]) -> bool:
    return all(len(graph.adj[node]) == 3 for node in graph.nodes - leaves)


def numeric_lengths(
    empirical_index: Mapping[tuple[tuple[str, ...], tuple[str, ...]], tuple[str, str]]
) -> dict[tuple[tuple[str, ...], tuple[str, ...]], Decimal]:
    return {
        split: Decimal(index) / Decimal(10)
        for index, split in enumerate(sorted(empirical_index, key=split_key), start=1)
    }


def decimal_text(value: Decimal | None) -> str:
    if value is None:
        return "NA"
    text = format(value, "f")
    return text.rstrip("0").rstrip(".") if "." in text else text


def primitive_output(reference_state: str, present: bool) -> str:
    if reference_state == "structural":
        return "NA_struct"
    if reference_state == "fused":
        return "NA_fuse"
    if reference_state != "eligible":
        raise ValueError(reference_state)
    return "Mapped" if present else "NA_topo"


@dataclass(frozen=True)
class GraphCompositeState:
    state: str
    components: tuple[frozenset[str], ...]
    saturated: bool


def graph_composite_state(
    deleted: frozenset[str], fibers: tuple[frozenset[str], ...], coordinate: frozenset[str]
) -> GraphCompositeState:
    components = tuple(sorted((fiber for fiber in fibers if fiber <= coordinate), key=coord_sort))
    deleted_free = not bool(set(coordinate) & set(deleted))
    whole = all(not (set(fiber) & set(coordinate)) or fiber <= coordinate for fiber in fibers)
    saturated = deleted_free and whole
    if coordinate in fibers:
        state = "active"
    elif saturated and len(components) >= 2:
        state = "decomposable"
    else:
        state = "unavailable"
    return GraphCompositeState(state, components, saturated)


def graph_composite_recovery(
    state: GraphCompositeState,
    component_splits: tuple[tuple[tuple[str, ...], tuple[str, ...]], ...],
    empirical_index: Mapping[tuple[tuple[str, ...], tuple[str, ...]], tuple[str, str]],
    lengths: Mapping[tuple[tuple[str, ...], tuple[str, ...]], Decimal],
) -> tuple[str, str, Decimal | None]:
    if state.state == "active" and len(component_splits) == 1 and component_splits[0] in empirical_index:
        return "recovered_active", "single_edge_lookup", lengths[component_splits[0]]
    if state.state == "decomposable" and all(split in empirical_index for split in component_splits):
        return (
            "recovered_decomposable",
            "selected_component_edge_sum",
            sum((lengths[split] for split in component_splits), Decimal(0)),
        )
    return "unrecovered", "no_numeric_value", None


@dataclass(frozen=True)
class SplitCompositeEvaluation:
    reference_state: str
    components: tuple[frozenset[str], ...]
    component_splits: tuple[tuple[tuple[str, ...], tuple[str, ...]], ...]
    saturated: bool
    recovery_state: str
    construction_mode: str
    numeric_value: Decimal | None


def evaluate_split_composite(
    structural: frozenset[str],
    groups: tuple[frozenset[str], ...],
    group_to_split: Mapping[frozenset[str], tuple[tuple[str, ...], tuple[str, ...]]],
    coordinate: frozenset[str],
    empirical_index: Mapping[tuple[tuple[str, ...], tuple[str, ...]], tuple[str, str]],
    lengths: Mapping[tuple[tuple[str, ...], tuple[str, ...]], Decimal],
) -> SplitCompositeEvaluation:
    """Evaluate a composite only through split groups and empirical split queries."""
    coordinate_values = set(coordinate)
    selected: list[frozenset[str]] = []
    covered: set[str] = set()
    partial_group = False
    for group in groups:
        overlap = coordinate_values.intersection(group)
        if not overlap:
            continue
        if overlap == set(group):
            selected.append(group)
            covered.update(group)
        else:
            partial_group = True
    components = tuple(sorted(selected, key=coord_sort))
    saturated = (
        not coordinate_values.intersection(structural)
        and not partial_group
        and covered == coordinate_values
    )
    direct_group_match = any(group == coordinate for group in groups)
    if direct_group_match:
        reference_state = "active"
    elif saturated and len(components) > 1:
        reference_state = "decomposable"
    else:
        reference_state = "unavailable"

    component_splits = tuple(group_to_split[group] for group in components)
    recovery_state = "unrecovered"
    construction_mode = "no_numeric_value"
    numeric_value: Decimal | None = None
    if reference_state == "active":
        if len(component_splits) == 1:
            selected_split = component_splits[0]
            empirical_edge = empirical_index.get(selected_split)
            if empirical_edge is not None and selected_split in lengths:
                recovery_state = "recovered_active"
                construction_mode = "single_edge_lookup"
                numeric_value = lengths[selected_split]
    elif reference_state == "decomposable":
        selected_values: list[Decimal] = []
        complete = True
        for selected_split in component_splits:
            empirical_edge = empirical_index.get(selected_split)
            if empirical_edge is None or selected_split not in lengths:
                complete = False
                break
            selected_values.append(lengths[selected_split])
        if complete:
            recovery_state = "recovered_decomposable"
            construction_mode = "selected_component_edge_sum"
            numeric_value = sum(selected_values, Decimal(0))
    return SplitCompositeEvaluation(
        reference_state,
        components,
        component_splits,
        saturated,
        recovery_state,
        construction_mode,
        numeric_value,
    )


@dataclass(frozen=True)
class BoundaryLookup:
    member_set: frozenset[str]
    reference_state: str
    lookup_split: tuple[tuple[str, ...], tuple[str, ...]] | None
    recovery_state: str
    construction_mode: str
    numeric_value: Decimal | None


def two_leaf_empirical_graph(left: str, right: str) -> Graph:
    empirical = Graph()
    empirical.add_edge(left, right, "two_leaf_empirical_edge")
    return empirical


def graph_two_taxon_lookup(
    reference: Graph,
    coordinate_to_edge: Mapping[str, tuple[str, str]],
    retained: frozenset[str],
    restriction: Restriction,
    empirical_index: Mapping[tuple[tuple[str, ...], tuple[str, ...]], tuple[str, str]],
    lengths: Mapping[tuple[tuple[str, ...], tuple[str, ...]], Decimal],
) -> BoundaryLookup:
    if len(restriction.fibers) != 1:
        return BoundaryLookup(
            frozenset(), "unavailable", None, "unrecovered", "no_numeric_value", None
        )
    member_set = restriction.fibers[0]
    state = graph_composite_state(restriction.deleted, restriction.fibers, member_set)
    representative = min(member_set, key=COORD_INDEX.__getitem__)
    lookup_split = projected_split(reference, coordinate_to_edge[representative], set(retained))
    component_splits = (lookup_split,) if lookup_split is not None else ()
    recovery, mode, value = graph_composite_recovery(
        state, component_splits, empirical_index, lengths
    )
    return BoundaryLookup(member_set, state.state, lookup_split, recovery, mode, value)


def split_two_taxon_lookup(
    restriction: SplitRestriction,
    empirical_index: Mapping[tuple[tuple[str, ...], tuple[str, ...]], tuple[str, str]],
    lengths: Mapping[tuple[tuple[str, ...], tuple[str, ...]], Decimal],
) -> BoundaryLookup:
    if len(restriction.groups) != 1:
        return BoundaryLookup(
            frozenset(), "unavailable", None, "unrecovered", "no_numeric_value", None
        )
    member_set = restriction.groups[0]
    lookup_split = restriction.group_to_split.get(member_set)
    reference_state = "active" if lookup_split is not None else "unavailable"
    recovery_state = "unrecovered"
    construction_mode = "no_numeric_value"
    numeric_value: Decimal | None = None
    if reference_state == "active" and lookup_split is not None:
        empirical_edge = empirical_index.get(lookup_split)
        if empirical_edge is not None and lookup_split in lengths:
            recovery_state = "recovered_active"
            construction_mode = "single_edge_lookup"
            numeric_value = lengths[lookup_split]
    return BoundaryLookup(
        member_set,
        reference_state,
        lookup_split,
        recovery_state,
        construction_mode,
        numeric_value,
    )


def write_tsv(path: Path, rows: Sequence[Mapping[str, object]], fields: Sequence[str]) -> None:
    path.parent.mkdir(parents=True, exist_ok=True)
    with path.open("w", encoding="utf-8", newline="") as handle:
        writer = csv.DictWriter(handle, fieldnames=fields, delimiter="\t", lineterminator="\n")
        writer.writeheader()
        for row in rows:
            writer.writerow({field: row.get(field, "") for field in fields})


def add_discrepancy(
    rows: list[dict[str, str]], domain: str, case_id: str, coordinate_type: str,
    coordinate_id: str, check: str, expected: object, observed: object, detail: str = ""
) -> None:
    rows.append(
        {
            "discrepancy_id": f"D{len(rows)+1:05d}",
            "domain": domain,
            "case_id": case_id,
            "coordinate_type": coordinate_type,
            "coordinate_id": coordinate_id,
            "check": check,
            "expected": str(expected),
            "observed": str(observed),
            "detail": detail,
        }
    )


def result_row(
    invariant_id: str, invariant: str, expected: object, observed: object, passed: bool, scope: str
) -> dict[str, str]:
    return {
        "invariant_id": invariant_id,
        "invariant": invariant,
        "scope": scope,
        "expected": str(expected),
        "observed": str(observed),
        "pass": str(passed).lower(),
    }


def file_sha256(path: Path) -> str:
    digest = hashlib.sha256()
    with path.open("rb") as handle:
        for chunk in iter(lambda: handle.read(1024 * 1024), b""):
            digest.update(chunk)
    return digest.hexdigest()


def main() -> int:
    parser = argparse.ArgumentParser()
    parser.add_argument("--output-dir", required=True)
    args = parser.parse_args()
    root = Path(args.output_dir)
    root.mkdir(parents=True, exist_ok=False)
    machine = root / "outputs"
    machine.mkdir()

    reference, coordinate_to_edge = reference_graph()
    discrepancies: list[dict[str, str]] = []
    restrictions: dict[frozenset[str], Restriction] = {}
    split_restrictions: dict[frozenset[str], SplitRestriction] = {}
    restriction_rows: list[dict[str, str]] = []
    order_rows: list[dict[str, str]] = []
    composite_axis: set[frozenset[str]] = set()
    endpoint_collapse_count = 0
    recurrence_expected_counts = {
        size: unrooted_topology_count_recurrence(size) for size in range(3, 7)
    }
    for size, declared in EXPECTED_TOPOLOGY_COUNTS.items():
        if recurrence_expected_counts[size] != declared:
            add_discrepancy(
                discrepancies,
                "topology_recurrence",
                f"n{size}",
                "topology",
                "all",
                "recurrence_vs_frozen_protocol",
                declared,
                recurrence_expected_counts[size],
            )

    retained_sets = [
        frozenset(values)
        for size in range(3, 7)
        for values in itertools.combinations(TAXA, size)
    ]
    for retained in retained_sets:
        retained_id = "R_" + taxa_key(retained)
        graph_result = graph_restriction(reference, coordinate_to_edge, retained)
        split_result = split_restriction(reference, coordinate_to_edge, retained)
        restrictions[retained] = graph_result
        split_restrictions[retained] = split_result
        composite_axis.update(fiber for fiber in graph_result.fibers if len(fiber) >= 2)
        if graph_result.deleted != split_result.structural:
            add_discrepancy(
                discrepancies, "restriction", retained_id, "all", "all", "deleted_vs_empty_projected",
                member_key(graph_result.deleted), member_key(split_result.structural)
            )
        if set(graph_result.fibers) != set(split_result.groups):
            add_discrepancy(
                discrepancies, "restriction", retained_id, "all", "all", "fiber_split_equivalence",
                member_sets_text(graph_result.fibers), member_sets_text(split_result.groups)
            )
        ascending = suppression_partition(reference, coordinate_to_edge, set(graph_result.k_edges), False)
        descending = suppression_partition(reference, coordinate_to_edge, set(graph_result.k_edges), True)
        for name, observed in (("ascending", ascending), ("descending", descending)):
            passed = observed == graph_result.fibers
            order_rows.append(
                {
                    "trial_id": f"{retained_id}_SUP_{name}",
                    "retained_id": retained_id,
                    "trial_type": "suppression_order",
                    "order": name,
                    "expected_k_edges": member_key(reference.labels[edge] for edge in graph_result.k_edges),
                    "observed_k_edges": member_key(reference.labels[edge] for edge in graph_result.k_edges),
                    "expected_fibers": member_sets_text(graph_result.fibers),
                    "observed_fibers": member_sets_text(observed),
                    "k_edges_match": "true",
                    "fibers_match": str(passed).lower(),
                    "pass": str(passed).lower(),
                }
            )
            if not passed:
                add_discrepancy(
                    discrepancies, "suppression", retained_id, "all", "all", "suppression_order",
                    member_sets_text(graph_result.fibers), member_sets_text(observed), name
                )
        for index, order in enumerate(itertools.permutations(sorted(retained)), start=1):
            observed_k = k_edges_from_order(reference, order)
            observed_restriction = restriction_from_k_edges(
                reference, coordinate_to_edge, retained, observed_k
            )
            k_edges_match = observed_k == set(graph_result.k_edges)
            fibers_match = observed_restriction.fibers == graph_result.fibers
            passed = k_edges_match and fibers_match
            order_rows.append(
                {
                    "trial_id": f"{retained_id}_ORD_{index:04d}",
                    "retained_id": retained_id,
                    "trial_type": "construction_order",
                    "order": ",".join(order),
                    "expected_k_edges": member_key(reference.labels[edge] for edge in graph_result.k_edges),
                    "observed_k_edges": member_key(reference.labels[edge] for edge in observed_k),
                    "expected_fibers": member_sets_text(graph_result.fibers),
                    "observed_fibers": member_sets_text(observed_restriction.fibers),
                    "k_edges_match": str(k_edges_match).lower(),
                    "fibers_match": str(fibers_match).lower(),
                    "pass": str(passed).lower(),
                }
            )
            if not k_edges_match:
                add_discrepancy(
                    discrepancies, "construction", retained_id, "all", "all", "construction_order_k",
                    member_key(reference.labels[edge] for edge in graph_result.k_edges),
                    member_key(reference.labels[edge] for edge in observed_k),
                    ",".join(order)
                )
            if not fibers_match:
                add_discrepancy(
                    discrepancies, "construction", retained_id, "all", "all", "construction_order_fibers",
                    member_sets_text(graph_result.fibers), member_sets_text(observed_restriction.fibers),
                    ",".join(order)
                )
        path_verified = True
        for fiber in graph_result.fibers:
            edges = {coordinate_to_edge[coordinate] for coordinate in fiber}
            degrees: Counter[str] = Counter(node for edge in edges for node in edge)
            adjacency = {edge: set() for edge in edges}
            for first, second in itertools.combinations(edges, 2):
                if set(first) & set(second):
                    adjacency[first].add(second)
                    adjacency[second].add(first)
            connected = len(connected_edge_components(edges, adjacency)) == 1 if edges else True
            if not connected or any(value > 2 for value in degrees.values()):
                path_verified = False
        for coordinate in TAXA:
            fiber = graph_result.coordinate_to_fiber[coordinate]
            if coordinate in retained and fiber is not None and len(fiber) >= 2:
                endpoint_collapse_count += 1
        restriction_rows.append(
            {
                "retained_id": retained_id,
                "retained_taxa": ",".join(sorted(retained)),
                "retained_count": str(len(retained)),
                "k_edges": member_key(reference.labels[edge] for edge in graph_result.k_edges),
                "deleted_edges": member_key(graph_result.deleted),
                "graph_fibers": member_sets_text(graph_result.fibers),
                "split_groups": member_sets_text(split_result.groups),
                "fiber_split_equivalent": str(set(graph_result.fibers) == set(split_result.groups)).lower(),
                "fiber_paths_verified": str(path_verified).lower(),
                "suppression_ascending": member_sets_text(ascending),
                "suppression_descending": member_sets_text(descending),
                "suppression_order_independent": str(ascending == descending == graph_result.fibers).lower(),
            }
        )

    if composite_axis != EXPECTED_COMPOSITE_AXIS:
        add_discrepancy(
            discrepancies, "axis", "all", "composite", "B_star", "composite_axis",
            member_sets_text(EXPECTED_COMPOSITE_AXIS), member_sets_text(composite_axis)
        )
    composite_axis_ordered = tuple(sorted(composite_axis, key=coord_sort))

    topology_cache: dict[tuple[str, ...], dict[tuple[str, ...], Graph]] = {}
    topology_counts: dict[int, int] = {}
    binary_counts: dict[int, int] = {}
    case_rows: list[dict[str, str]] = []
    primitive_rows: list[dict[str, str]] = []
    composite_rows: list[dict[str, str]] = []

    for retained in retained_sets:
        labels = tuple(sorted(retained))
        if labels not in topology_cache:
            topology_cache[labels] = all_normalized_topologies(labels)
        topologies = topology_cache[labels]
        topology_counts[len(labels)] = len(topologies)
        binary_count = sum(topology_is_binary(graph, set(labels)) for graph in topologies.values())
        binary_counts[len(labels)] = binary_count
        if len(topologies) != recurrence_expected_counts[len(labels)]:
            add_discrepancy(
                discrepancies, "topology", "R_" + taxa_key(retained), "topology", "all",
                "topology_count", recurrence_expected_counts[len(labels)], len(topologies)
            )
        if binary_count != EXPECTED_BINARY_COUNTS[len(labels)]:
            add_discrepancy(
                discrepancies, "topology", "R_" + taxa_key(retained), "topology", "binary",
                "binary_topology_count", EXPECTED_BINARY_COUNTS[len(labels)], binary_count
            )
        graph_result = restrictions[retained]
        split_result = split_restrictions[retained]
        for topology_index, signature in enumerate(sorted(topologies), start=1):
            empirical = topologies[signature]
            empirical_index = empirical_split_index(empirical, set(retained))
            lengths = numeric_lengths(empirical_index)
            resolution = "binary" if topology_is_binary(empirical, set(retained)) else "multifurcating"
            case_id = f"M_R{taxa_key(retained)}_T{topology_index:03d}"
            internal_split_count = sum(
                1 for split in empirical_index if len(split[0]) >= 2 and len(split[1]) >= 2
            )
            case_rows.append(
                {
                    "case_id": case_id,
                    "retained_id": "R_" + taxa_key(retained),
                    "retained_taxa": ",".join(sorted(retained)),
                    "retained_count": str(len(retained)),
                    "topology_id": f"U{len(retained)}_{topology_index:03d}",
                    "resolution_class": resolution,
                    "empirical_edge_count": str(len(empirical.edges)),
                    "empirical_internal_split_count": str(internal_split_count),
                    "topology_signature": ";".join(signature),
                }
            )
            for coordinate in COORDS:
                graph_ref = graph_result.reference_state[coordinate]
                split_ref = split_result.reference_state[coordinate]
                graph_fiber = graph_result.coordinate_to_fiber[coordinate]
                split_group = split_result.coordinate_to_group[coordinate]
                split = (
                    projected_split(reference, coordinate_to_edge[coordinate], set(retained))
                    if graph_ref != "structural"
                    else None
                )
                present = split in empirical_index if split is not None else False
                graph_output = primitive_output(graph_ref, present)
                split_output = primitive_output(split_ref, present)
                mode = "single_edge_lookup" if graph_output == "Mapped" else "no_numeric_value"
                numeric = lengths[split] if graph_output == "Mapped" and split is not None else None
                if graph_ref != split_ref:
                    add_discrepancy(
                        discrepancies, "main", case_id, "primitive", coordinate, "reference_state",
                        graph_ref, split_ref
                    )
                if graph_fiber != split_group:
                    add_discrepancy(
                        discrepancies, "main", case_id, "primitive", coordinate, "fiber_group",
                        member_key(graph_fiber or []), member_key(split_group or [])
                    )
                if graph_output != split_output:
                    add_discrepancy(
                        discrepancies, "main", case_id, "primitive", coordinate, "output_state",
                        graph_output, split_output
                    )
                if coordinate in TAXA and coordinate in retained and graph_output in {"NA_struct", "NA_topo"}:
                    add_discrepancy(
                        discrepancies, "main", case_id, "primitive", coordinate,
                        "retained_terminal_boundary", "Mapped or NA_fuse", graph_output
                    )
                primitive_rows.append(
                    {
                        "cell_id": f"{case_id}|P|{coordinate}",
                        "case_id": case_id,
                        "coordinate": coordinate,
                        "graph_reference_state": graph_ref,
                        "split_reference_state": split_ref,
                        "graph_fiber": member_key(graph_fiber or [] ) if graph_fiber else "bottom",
                        "split_group": member_key(split_group or []) if split_group else "bottom",
                        "projected_split": split_key(split) if split is not None else "NA",
                        "empirical_split_present": str(present).lower() if split is not None else "NA",
                        "graph_output_state": graph_output,
                        "split_output_state": split_output,
                        "construction_mode": mode,
                        "numeric_value": decimal_text(numeric),
                    }
                )
            for coordinate in composite_axis_ordered:
                graph_state = graph_composite_state(
                    graph_result.deleted, graph_result.fibers, coordinate
                )
                graph_splits = tuple(
                    projected_split(
                        reference,
                        coordinate_to_edge[min(component, key=COORD_INDEX.__getitem__)],
                        set(retained),
                    )
                    for component in graph_state.components
                )
                if any(value is None for value in graph_splits):
                    raise AssertionError("nondeleted graph component has empty projected split")
                graph_splits_typed = tuple(value for value in graph_splits if value is not None)
                graph_recovery, graph_mode, graph_numeric = graph_composite_recovery(
                    graph_state, graph_splits_typed, empirical_index, lengths
                )
                split_evaluation = evaluate_split_composite(
                    split_result.structural,
                    split_result.groups,
                    split_result.group_to_split,
                    coordinate,
                    empirical_index,
                    lengths,
                )
                coordinate_id = member_key(coordinate)
                if graph_state.state != split_evaluation.reference_state:
                    add_discrepancy(
                        discrepancies, "main", case_id, "composite", coordinate_id,
                        "composite_reference", graph_state.state, split_evaluation.reference_state
                    )
                if graph_state.components != split_evaluation.components:
                    add_discrepancy(
                        discrepancies, "main", case_id, "composite", coordinate_id,
                        "composite_components", member_sets_text(graph_state.components),
                        member_sets_text(split_evaluation.components)
                    )
                if graph_state.saturated != split_evaluation.saturated:
                    add_discrepancy(
                        discrepancies, "main", case_id, "composite", coordinate_id,
                        "composite_saturation", graph_state.saturated, split_evaluation.saturated
                    )
                if graph_splits_typed != split_evaluation.component_splits:
                    add_discrepancy(
                        discrepancies, "main", case_id, "composite", coordinate_id,
                        "component_splits", ";".join(map(split_key, graph_splits_typed)),
                        ";".join(map(split_key, split_evaluation.component_splits))
                    )
                if (graph_recovery, graph_mode, graph_numeric) != (
                    split_evaluation.recovery_state,
                    split_evaluation.construction_mode,
                    split_evaluation.numeric_value,
                ):
                    add_discrepancy(
                        discrepancies, "main", case_id, "composite", coordinate_id,
                        "composite_recovery_numeric",
                        (graph_recovery, graph_mode, decimal_text(graph_numeric)),
                        (
                            split_evaluation.recovery_state,
                            split_evaluation.construction_mode,
                            decimal_text(split_evaluation.numeric_value),
                        )
                    )
                composite_rows.append(
                    {
                        "cell_id": f"{case_id}|C|{coordinate_id}",
                        "case_id": case_id,
                        "coordinate_member_set": coordinate_id,
                        "graph_reference_state": graph_state.state,
                        "split_reference_state": split_evaluation.reference_state,
                        "graph_saturated": str(graph_state.saturated).lower(),
                        "split_saturated": str(split_evaluation.saturated).lower(),
                        "graph_components": member_sets_text(graph_state.components),
                        "split_components": member_sets_text(split_evaluation.components),
                        "graph_component_splits": ";".join(map(split_key, graph_splits_typed)) or "NA",
                        "split_component_splits": ";".join(
                            map(split_key, split_evaluation.component_splits)
                        ) or "NA",
                        "graph_recovery_state": graph_recovery,
                        "split_recovery_state": split_evaluation.recovery_state,
                        "graph_construction_mode": graph_mode,
                        "split_construction_mode": split_evaluation.construction_mode,
                        "graph_numeric_value": decimal_text(graph_numeric),
                        "split_numeric_value": decimal_text(split_evaluation.numeric_value),
                    }
                )

    boundary_rows: list[dict[str, str]] = []
    boundary_case_ids: set[str] = set()
    for retained_size in (2, 1):
        for retained_tuple in itertools.combinations(TAXA, retained_size):
            retained = frozenset(retained_tuple)
            boundary_class = "two_taxon" if retained_size == 2 else "one_taxon"
            case_id = f"B_{retained_size}_{taxa_key(retained)}"
            boundary_case_ids.add(case_id)
            graph_result = graph_restriction(reference, coordinate_to_edge, retained)
            split_result = split_restriction(reference, coordinate_to_edge, retained)
            boundary_empirical_index: dict[
                tuple[tuple[str, ...], tuple[str, ...]], tuple[str, str]
            ] = {}
            boundary_lengths: dict[
                tuple[tuple[str, ...], tuple[str, ...]], Decimal
            ] = {}
            boundary_topology_signature = "NA"
            boundary_empirical_edge_count = "NA"
            if retained_size == 2:
                boundary_empirical = two_leaf_empirical_graph(*retained_tuple)
                boundary_empirical_index = empirical_split_index(
                    boundary_empirical, set(retained)
                )
                if len(boundary_empirical_index) != 1:
                    add_discrepancy(
                        discrepancies,
                        "boundary",
                        case_id,
                        "empirical",
                        "two_leaf",
                        "empirical_split_count",
                        1,
                        len(boundary_empirical_index),
                    )
                boundary_lengths = {
                    split: Decimal("1") for split in boundary_empirical_index
                }
                boundary_topology_signature = ";".join(
                    sorted(split_key(split) for split in boundary_empirical_index)
                )
                boundary_empirical_edge_count = str(len(boundary_empirical.edges))
            if set(graph_result.fibers) != set(split_result.groups):
                add_discrepancy(
                    discrepancies, "boundary", case_id, "all", "all", "fiber_split_equivalence",
                    member_sets_text(graph_result.fibers), member_sets_text(split_result.groups)
                )
            for coordinate in COORDS:
                graph_ref = graph_result.reference_state[coordinate]
                split_ref = split_result.reference_state[coordinate]
                output = "NA_struct" if graph_ref == "structural" else "NA_fuse"
                expected = "NA_struct" if retained_size == 1 or coordinate in graph_result.deleted else "NA_fuse"
                passed = graph_ref == split_ref and output == expected
                boundary_rows.append(
                    {
                        "case_id": case_id,
                        "boundary_class": boundary_class,
                        "retained_taxa": ",".join(sorted(retained)),
                        "row_type": "primitive",
                        "coordinate_id": coordinate,
                        "graph_reference_state": graph_ref,
                        "split_reference_state": split_ref,
                        "output_state": output,
                        "graph_recovery_state": "NA",
                        "split_recovery_state": "NA",
                        "graph_construction_mode": "no_numeric_value",
                        "split_construction_mode": "no_numeric_value",
                        "graph_numeric_value": "NA",
                        "split_numeric_value": "NA",
                        "path_members": member_key(next(iter(graph_result.fibers), frozenset())),
                        "empirical_topology_signature": "NA",
                        "empirical_edge_count": "NA",
                        "graph_lookup_split": "NA",
                        "split_lookup_split": "NA",
                        "declared_length_rule": "NA",
                        "pass": str(passed).lower(),
                    }
                )
                if not passed:
                    add_discrepancy(
                        discrepancies, "boundary", case_id, "primitive", coordinate, "boundary_state",
                        expected, output
                    )
            if retained_size == 2:
                graph_lookup = graph_two_taxon_lookup(
                    reference,
                    coordinate_to_edge,
                    retained,
                    graph_result,
                    boundary_empirical_index,
                    boundary_lengths,
                )
                split_lookup = split_two_taxon_lookup(
                    split_result, boundary_empirical_index, boundary_lengths
                )
                if graph_lookup.member_set != split_lookup.member_set:
                    add_discrepancy(
                        discrepancies,
                        "boundary",
                        case_id,
                        "composite",
                        "path",
                        "two_taxon_member_set",
                        member_key(graph_lookup.member_set),
                        member_key(split_lookup.member_set),
                    )
                if graph_lookup.reference_state != split_lookup.reference_state:
                    add_discrepancy(
                        discrepancies,
                        "boundary",
                        case_id,
                        "composite",
                        "path",
                        "two_taxon_reference_state",
                        graph_lookup.reference_state,
                        split_lookup.reference_state,
                    )
                if graph_lookup.lookup_split != split_lookup.lookup_split:
                    add_discrepancy(
                        discrepancies,
                        "boundary",
                        case_id,
                        "composite",
                        "path",
                        "two_taxon_lookup_split",
                        split_key(graph_lookup.lookup_split) if graph_lookup.lookup_split else "NA",
                        split_key(split_lookup.lookup_split) if split_lookup.lookup_split else "NA",
                    )
                graph_lookup_tuple = (
                    graph_lookup.recovery_state,
                    graph_lookup.construction_mode,
                    graph_lookup.numeric_value,
                )
                split_lookup_tuple = (
                    split_lookup.recovery_state,
                    split_lookup.construction_mode,
                    split_lookup.numeric_value,
                )
                expected_lookup_tuple = (
                    "recovered_active",
                    "single_edge_lookup",
                    Decimal("1"),
                )
                if graph_lookup_tuple != split_lookup_tuple:
                    add_discrepancy(
                        discrepancies,
                        "boundary",
                        case_id,
                        "composite",
                        "path",
                        "two_taxon_recovery_numeric",
                        (
                            graph_lookup.recovery_state,
                            graph_lookup.construction_mode,
                            decimal_text(graph_lookup.numeric_value),
                        ),
                        (
                            split_lookup.recovery_state,
                            split_lookup.construction_mode,
                            decimal_text(split_lookup.numeric_value),
                        ),
                    )
                boundary_passed = (
                    len(graph_result.fibers) == 1
                    and len(split_result.groups) == 1
                    and graph_lookup.member_set == split_lookup.member_set
                    and graph_lookup.reference_state == split_lookup.reference_state == "active"
                    and graph_lookup.lookup_split == split_lookup.lookup_split
                    and graph_lookup_tuple == split_lookup_tuple == expected_lookup_tuple
                    and len(boundary_empirical_index) == 1
                    and boundary_empirical_edge_count == "1"
                )
                if not boundary_passed and graph_lookup_tuple == split_lookup_tuple:
                    add_discrepancy(
                        discrepancies,
                        "boundary",
                        case_id,
                        "composite",
                        "path",
                        "two_taxon_expected_lookup",
                        ("active", "recovered_active", "single_edge_lookup", "1"),
                        (
                            graph_lookup.reference_state,
                            graph_lookup.recovery_state,
                            graph_lookup.construction_mode,
                            decimal_text(graph_lookup.numeric_value),
                        ),
                    )
                boundary_rows.append(
                    {
                        "case_id": case_id,
                        "boundary_class": boundary_class,
                        "retained_taxa": ",".join(sorted(retained)),
                        "row_type": "composite",
                        "coordinate_id": member_key(graph_lookup.member_set),
                        "graph_reference_state": graph_lookup.reference_state,
                        "split_reference_state": split_lookup.reference_state,
                        "output_state": "NA",
                        "graph_recovery_state": graph_lookup.recovery_state,
                        "split_recovery_state": split_lookup.recovery_state,
                        "graph_construction_mode": graph_lookup.construction_mode,
                        "split_construction_mode": split_lookup.construction_mode,
                        "graph_numeric_value": decimal_text(graph_lookup.numeric_value),
                        "split_numeric_value": decimal_text(split_lookup.numeric_value),
                        "path_members": member_key(graph_lookup.member_set),
                        "empirical_topology_signature": boundary_topology_signature,
                        "empirical_edge_count": boundary_empirical_edge_count,
                        "graph_lookup_split": split_key(graph_lookup.lookup_split)
                        if graph_lookup.lookup_split
                        else "NA",
                        "split_lookup_split": split_key(split_lookup.lookup_split)
                        if split_lookup.lookup_split
                        else "NA",
                        "declared_length_rule": 'single empirical edge assigned Decimal("1")',
                        "pass": str(boundary_passed).lower(),
                    }
                )

    main_case_count = len(case_rows)
    binary_case_count = sum(row["resolution_class"] == "binary" for row in case_rows)
    topology_count_by_size = {
        size: len(all_normalized_topologies(tuple(TAXA[:size]))) for size in range(3, 7)
    }
    binary_count_by_size = {
        size: sum(
            topology_is_binary(graph, set(TAXA[:size]))
            for graph in all_normalized_topologies(tuple(TAXA[:size])).values()
        )
        for size in range(3, 7)
    }
    topology_recurrence_rows = []
    for size in range(3, 7):
        recurrence_count = recurrence_expected_counts[size]
        generated_count = topology_count_by_size[size]
        declared_count = EXPECTED_TOPOLOGY_COUNTS[size]
        recurrence_passed = recurrence_count == declared_count == generated_count
        topology_recurrence_rows.append(
            {
                "retained_taxa_count": size,
                "rooted_recurrence_leaf_count": size - 1,
                "declared_protocol_count": declared_count,
                "partition_recurrence_count": recurrence_count,
                "generated_topology_count": generated_count,
                "pass": str(recurrence_passed).lower(),
            }
        )
        if not recurrence_passed:
            add_discrepancy(
                discrepancies,
                "topology_recurrence",
                f"n{size}",
                "topology",
                "all",
                "partition_recurrence_reconciliation",
                (declared_count, recurrence_count),
                generated_count,
            )

    case_count_rows = [
        {"count_id": "retained_sets_main", "expected": 42, "observed": len(retained_sets)},
        *[
            {
                "count_id": f"topology_recurrence_k{size}",
                "expected": EXPECTED_TOPOLOGY_COUNTS[size],
                "observed": recurrence_expected_counts[size],
            }
            for size in range(3, 7)
        ],
        *[
            {
                "count_id": f"topologies_k{size}",
                "expected": recurrence_expected_counts[size],
                "observed": topology_count_by_size[size],
            }
            for size in range(3, 7)
        ],
        {"count_id": "main_cases_all", "expected": 472, "observed": main_case_count},
        {"count_id": "main_cases_binary_subset", "expected": 260, "observed": binary_case_count},
        {"count_id": "composite_axis", "expected": 21, "observed": len(composite_axis)},
        {"count_id": "primitive_main_rows", "expected": 4248, "observed": len(primitive_rows)},
        {"count_id": "composite_main_rows", "expected": 9912, "observed": len(composite_rows)},
        {"count_id": "main_categorical_rows", "expected": 14160, "observed": len(primitive_rows) + len(composite_rows)},
        {"count_id": "construction_order_trials", "expected": 1920, "observed": sum(row["trial_type"] == "construction_order" for row in order_rows)},
        {"count_id": "suppression_order_trials", "expected": 84, "observed": sum(row["trial_type"] == "suppression_order" for row in order_rows)},
        {"count_id": "boundary_cases", "expected": 21, "observed": len(boundary_case_ids)},
        {"count_id": "boundary_primitive_rows", "expected": 189, "observed": sum(row["row_type"] == "primitive" for row in boundary_rows)},
        {"count_id": "boundary_composite_rows", "expected": 15, "observed": sum(row["row_type"] == "composite" for row in boundary_rows)},
    ]
    for row in case_count_rows:
        row["pass"] = str(row["expected"] == row["observed"]).lower()
        if row["expected"] != row["observed"]:
            add_discrepancy(
                discrepancies, "count", "all", "all", row["count_id"], "case_count",
                row["expected"], row["observed"]
            )

    coverage_specs = [
        ("primitive_reference", "structural", sum(row["graph_reference_state"] == "structural" for row in primitive_rows)),
        ("primitive_reference", "fused", sum(row["graph_reference_state"] == "fused" for row in primitive_rows)),
        ("primitive_reference", "eligible", sum(row["graph_reference_state"] == "eligible" for row in primitive_rows)),
        *[("primitive_output", state, sum(row["graph_output_state"] == state for row in primitive_rows)) for state in ("Mapped", "NA_topo", "NA_fuse", "NA_struct")],
        *[("composite_reference", state, sum(row["graph_reference_state"] == state for row in composite_rows)) for state in ("active", "decomposable", "unavailable")],
        *[("composite_recovery", state, sum(row["graph_recovery_state"] == state for row in composite_rows)) for state in ("recovered_active", "recovered_decomposable", "unrecovered")],
        *[("construction_mode", state, sum(row["graph_construction_mode"] == state for row in composite_rows) + sum(row["construction_mode"] == state for row in primitive_rows)) for state in ("single_edge_lookup", "selected_component_edge_sum", "no_numeric_value")],
        ("topology_resolution", "binary", binary_case_count),
        ("topology_resolution", "multifurcating", main_case_count - binary_case_count),
        ("boundary_class", "two_taxon", sum(row["boundary_class"] == "two_taxon" and row["row_type"] == "primitive" for row in boundary_rows) // 9),
        ("boundary_class", "one_taxon", sum(row["boundary_class"] == "one_taxon" and row["row_type"] == "primitive" for row in boundary_rows) // 9),
        ("endpoint_collapse", "retained_terminal_in_non_singleton_fiber", endpoint_collapse_count),
        ("strengthened_check", "independent_split_composite_rows", len(composite_rows)),
        ("strengthened_check", "explicit_two_leaf_empirical_lookups", sum(row["boundary_class"] == "two_taxon" and row["row_type"] == "composite" for row in boundary_rows)),
        ("strengthened_check", "observed_k_fiber_trials", sum(row["trial_type"] == "construction_order" for row in order_rows)),
        ("strengthened_check", "partition_recurrence_sizes", sum(row["pass"] == "true" for row in topology_recurrence_rows)),
    ]
    coverage_rows = [
        {
            "coverage_family": family,
            "state_or_class": state,
            "observed_count": count,
            "required_minimum": 1,
            "pass": str(count >= 1).lower(),
        }
        for family, state, count in coverage_specs
    ]
    for row in coverage_rows:
        if int(row["observed_count"]) < 1:
            add_discrepancy(
                discrepancies, "coverage", "all", row["coverage_family"], row["state_or_class"],
                "state_coverage", 1, row["observed_count"]
            )

    invariant_rows = [
        result_row("I01", "ordered-anchor K and fibers derived from each observed K agree with all-pairs restriction", 1920, sum(row["pass"] == "true" and row["k_edges_match"] == "true" and row["fibers_match"] == "true" and row["trial_type"] == "construction_order" for row in order_rows), all(row["pass"] == "true" and row["k_edges_match"] == "true" and row["fibers_match"] == "true" for row in order_rows if row["trial_type"] == "construction_order"), "main retained sets"),
        result_row("I02", "degree-2 suppression order independence", 84, sum(row["pass"] == "true" and row["trial_type"] == "suppression_order" for row in order_rows), all(row["pass"] == "true" for row in order_rows if row["trial_type"] == "suppression_order"), "main retained sets"),
        result_row("I03", "deleted edges and fibers partition primitive axis", 42, sum((len(row["deleted_edges"].split("|")) if row["deleted_edges"] else 0) + sum(len(value.split("|")) for value in row["graph_fibers"].split(";") if value) == 9 for row in restriction_rows), all((len(row["deleted_edges"].split("|")) if row["deleted_edges"] else 0) + sum(len(value.split("|")) for value in row["graph_fibers"].split(";") if value) == 9 for row in restriction_rows), "main retained sets"),
        result_row("I04", "fiber-split equivalence", 42, sum(row["fiber_split_equivalent"] == "true" for row in restriction_rows), all(row["fiber_split_equivalent"] == "true" for row in restriction_rows), "main retained sets"),
        result_row("I05", "all fibers are connected reference paths", 42, sum(row["fiber_paths_verified"] == "true" for row in restriction_rows), all(row["fiber_paths_verified"] == "true" for row in restriction_rows), "main retained sets"),
        result_row("I06", "primitive graph/split agreement", 4248, sum(row["graph_output_state"] == row["split_output_state"] for row in primitive_rows), all(row["graph_output_state"] == row["split_output_state"] for row in primitive_rows), "main cells"),
        result_row("I07", "retained terminal never structural or topological NA", 0, sum(row["coordinate"] in TAXA and row["coordinate"] in case_rows_by_id(case_rows)[row["case_id"]] and row["graph_output_state"] in {"NA_struct", "NA_topo"} for row in primitive_rows), not any(row["coordinate"] in TAXA and row["coordinate"] in case_rows_by_id(case_rows)[row["case_id"]] and row["graph_output_state"] in {"NA_struct", "NA_topo"} for row in primitive_rows), "main terminal cells"),
        result_row("I08", "independent graph/split composite state, components, splits, recovery, mode, and value agreement", 9912, sum(row["graph_reference_state"] == row["split_reference_state"] and row["graph_saturated"] == row["split_saturated"] and row["graph_components"] == row["split_components"] and row["graph_component_splits"] == row["split_component_splits"] and row["graph_recovery_state"] == row["split_recovery_state"] and row["graph_construction_mode"] == row["split_construction_mode"] and row["graph_numeric_value"] == row["split_numeric_value"] for row in composite_rows), all(row["graph_reference_state"] == row["split_reference_state"] and row["graph_saturated"] == row["split_saturated"] and row["graph_components"] == row["split_components"] and row["graph_component_splits"] == row["split_component_splits"] and row["graph_recovery_state"] == row["split_recovery_state"] and row["graph_construction_mode"] == row["split_construction_mode"] and row["graph_numeric_value"] == row["split_numeric_value"] for row in composite_rows), "main composite cells"),
        result_row("I09", "cell IDs unique", len(primitive_rows) + len(composite_rows), len({row["cell_id"] for row in primitive_rows + composite_rows}), len({row["cell_id"] for row in primitive_rows + composite_rows}) == len(primitive_rows) + len(composite_rows), "main cells"),
        result_row("I10", "explicit normalized two-leaf empirical lookup agrees on state, recovery, mode, and value", 15, sum(row["boundary_class"] == "two_taxon" and row["row_type"] == "composite" and row["pass"] == "true" for row in boundary_rows), all(row["pass"] == "true" for row in boundary_rows if row["boundary_class"] == "two_taxon"), "boundary"),
        result_row("I11", "one-taxon boundary cases", 6, len({row["case_id"] for row in boundary_rows if row["boundary_class"] == "one_taxon" and row["pass"] == "true"}), all(row["pass"] == "true" for row in boundary_rows if row["boundary_class"] == "one_taxon"), "boundary"),
        result_row("I12", "all claimed states and classes covered", len(coverage_rows), sum(row["pass"] == "true" for row in coverage_rows), all(row["pass"] == "true" for row in coverage_rows), "coverage"),
        result_row("I13", "independent set-partition recurrence agrees with declared and generated topology counts", 4, sum(row["pass"] == "true" for row in topology_recurrence_rows), all(row["pass"] == "true" for row in topology_recurrence_rows), "topology counts"),
    ]
    for row in invariant_rows:
        if row["pass"] != "true":
            add_discrepancy(
                discrepancies, "invariant", "all", "all", row["invariant_id"], row["invariant"],
                row["expected"], row["observed"]
            )

    write_tsv(machine / "MAIN_CASES.tsv", case_rows, (
        "case_id", "retained_id", "retained_taxa", "retained_count", "topology_id",
        "resolution_class", "empirical_edge_count", "empirical_internal_split_count", "topology_signature"
    ))
    write_tsv(machine / "RESTRICTION_RESULTS.tsv", restriction_rows, (
        "retained_id", "retained_taxa", "retained_count", "k_edges", "deleted_edges",
        "graph_fibers", "split_groups", "fiber_split_equivalent", "fiber_paths_verified",
        "suppression_ascending", "suppression_descending", "suppression_order_independent"
    ))
    write_tsv(machine / "PRIMITIVE_RESULTS.tsv", primitive_rows, (
        "cell_id", "case_id", "coordinate", "graph_reference_state", "split_reference_state",
        "graph_fiber", "split_group", "projected_split", "empirical_split_present",
        "graph_output_state", "split_output_state", "construction_mode", "numeric_value"
    ))
    write_tsv(machine / "COMPOSITE_RESULTS.tsv", composite_rows, (
        "cell_id", "case_id", "coordinate_member_set", "graph_reference_state",
        "split_reference_state", "graph_saturated", "split_saturated", "graph_components", "split_components",
        "graph_component_splits", "split_component_splits", "graph_recovery_state", "split_recovery_state",
        "graph_construction_mode", "split_construction_mode", "graph_numeric_value", "split_numeric_value"
    ))
    write_tsv(machine / "ORDER_INVARIANCE.tsv", order_rows, (
        "trial_id", "retained_id", "trial_type", "order", "expected_k_edges", "observed_k_edges",
        "expected_fibers", "observed_fibers", "k_edges_match", "fibers_match", "pass"
    ))
    write_tsv(machine / "BOUNDARY_RESULTS.tsv", boundary_rows, (
        "case_id", "boundary_class", "retained_taxa", "row_type", "coordinate_id",
        "graph_reference_state", "split_reference_state", "output_state", "graph_recovery_state",
        "split_recovery_state", "graph_construction_mode", "split_construction_mode",
        "graph_numeric_value", "split_numeric_value", "path_members",
        "empirical_topology_signature", "empirical_edge_count", "graph_lookup_split",
        "split_lookup_split", "declared_length_rule", "pass"
    ))
    write_tsv(root / "TOPOLOGY_COUNT_RECURRENCE.tsv", topology_recurrence_rows, (
        "retained_taxa_count", "rooted_recurrence_leaf_count", "declared_protocol_count",
        "partition_recurrence_count", "generated_topology_count", "pass"
    ))
    write_tsv(root / "CASE_COUNT_RECONCILIATION.tsv", case_count_rows, (
        "count_id", "expected", "observed", "pass"
    ))
    write_tsv(root / "STATE_COVERAGE.tsv", coverage_rows, (
        "coverage_family", "state_or_class", "observed_count", "required_minimum", "pass"
    ))
    write_tsv(root / "INVARIANT_RESULTS.tsv", invariant_rows, (
        "invariant_id", "invariant", "scope", "expected", "observed", "pass"
    ))
    write_tsv(root / "DISCREPANCIES.tsv", discrepancies, (
        "discrepancy_id", "domain", "case_id", "coordinate_type", "coordinate_id", "check",
        "expected", "observed", "detail"
    ))

    summary = {
        "status": "PASS" if not discrepancies else "FAIL",
        "evidence_version": "v0.2",
        "reference_newick": "(A,(B,C),(D,(E,F)));",
        "retained_set_count": len(retained_sets),
        "main_case_count": main_case_count,
        "binary_case_count": binary_case_count,
        "multifurcating_case_count": main_case_count - binary_case_count,
        "primitive_main_rows": len(primitive_rows),
        "composite_axis_count": len(composite_axis),
        "composite_main_rows": len(composite_rows),
        "total_main_categorical_rows": len(primitive_rows) + len(composite_rows),
        "construction_order_trials": sum(row["trial_type"] == "construction_order" for row in order_rows),
        "suppression_order_trials": sum(row["trial_type"] == "suppression_order" for row in order_rows),
        "two_taxon_cases": 15,
        "one_taxon_cases": 6,
        "discrepancy_count": len(discrepancies),
        "partition_recurrence_counts": {
            str(size): recurrence_expected_counts[size] for size in range(3, 7)
        },
        "strengthened_checks": {
            "independent_split_composite_rows": len(composite_rows),
            "explicit_two_leaf_empirical_lookups": sum(
                row["boundary_class"] == "two_taxon" and row["row_type"] == "composite"
                for row in boundary_rows
            ),
            "observed_k_fiber_trials": sum(
                row["trial_type"] == "construction_order" for row in order_rows
            ),
            "topology_partition_recurrence_sizes": len(topology_recurrence_rows),
        },
        "state_counts": {
            "primitive_output": dict(sorted(Counter(row["graph_output_state"] for row in primitive_rows).items())),
            "composite_reference": dict(sorted(Counter(row["graph_reference_state"] for row in composite_rows).items())),
            "composite_recovery": dict(sorted(Counter(row["graph_recovery_state"] for row in composite_rows).items())),
        },
        "numeric_rule": "main domain: deterministic nonnegative Decimal length by sorted empirical split rank; two-taxon boundary: the explicit single empirical edge has Decimal(1); single-edge lookup or selected-component-edge sum",
    }
    (machine / "SUMMARY.json").write_text(json.dumps(summary, indent=2, sort_keys=True) + "\n", encoding="utf-8")

    hash_rows = []
    for path in sorted(p for p in root.rglob("*") if p.is_file() and p.name != "OUTPUT_SHA256SUMS.txt"):
        hash_rows.append(f"{file_sha256(path)}  {path.relative_to(root)}")
    (root / "OUTPUT_SHA256SUMS.txt").write_text("\n".join(hash_rows) + "\n", encoding="utf-8")
    print(json.dumps(summary, indent=2, sort_keys=True))
    return 0 if not discrepancies else 1


def case_rows_by_id(case_rows: Sequence[Mapping[str, str]]) -> dict[str, set[str]]:
    return {
        row["case_id"]: set(value for value in row["retained_taxa"].split(",") if value)
        for row in case_rows
    }


if __name__ == "__main__":
    raise SystemExit(main())
```
