## Supplementary Computational Evidence for "A Unique Graph-Theoretic Truth Table for Cross-Gene Branch Identity": static_independence_check.py

```
#!/usr/bin/env python3
"""AST-based checks for the four v0.2 implementation boundaries."""

from __future__ import annotations

import argparse
import ast
import csv
import json
from pathlib import Path


def function_map(tree: ast.AST) -> dict[str, ast.FunctionDef]:
    return {
        node.name: node
        for node in ast.walk(tree)
        if isinstance(node, ast.FunctionDef)
    }


def called_names(node: ast.AST) -> set[str]:
    result: set[str] = set()
    for value in ast.walk(node):
        if not isinstance(value, ast.Call):
            continue
        if isinstance(value.func, ast.Name):
            result.add(value.func.id)
        elif isinstance(value.func, ast.Attribute):
            result.add(value.func.attr)
    return result


def check_row(check_id: str, passed: bool, expected: str, observed: str, detail: str) -> dict[str, str]:
    return {
        "check_id": check_id,
        "status": "PASS" if passed else "FAIL",
        "expected": expected,
        "observed": observed,
        "detail": detail,
    }


def main() -> int:
    parser = argparse.ArgumentParser()
    parser.add_argument("validation_script")
    parser.add_argument("output_dir")
    args = parser.parse_args()
    script = Path(args.validation_script)
    output = Path(args.output_dir)
    output.mkdir(parents=True, exist_ok=False)
    tree = ast.parse(script.read_text(encoding="utf-8"), filename=str(script))
    functions = function_map(tree)
    required = {
        "graph_composite_state",
        "graph_composite_recovery",
        "evaluate_split_composite",
        "graph_two_taxon_lookup",
        "split_two_taxon_lookup",
        "restriction_from_k_edges",
        "graph_restriction",
        "labelled_set_partitions",
        "rooted_partition_tree_count",
        "unrooted_topology_count_recurrence",
        "all_normalized_topologies",
        "main",
    }
    missing = sorted(required - set(functions))
    rows = [
        check_row("S01_required_functions", not missing, "all present", ",".join(missing) or "all present", "")
    ]
    if not missing:
        split_calls = called_names(functions["evaluate_split_composite"])
        prohibited_composite = {
            "graph_composite_state",
            "graph_composite_recovery",
            "graph_two_taxon_lookup",
            "composite_state",
            "composite_recovery",
        }
        overlap = sorted(split_calls & prohibited_composite)
        rows.append(
            check_row(
                "S02_split_composite_no_graph_calls",
                not overlap,
                "no graph composite calls",
                ",".join(overlap) or "none",
                "split-side composite evaluator is a separate function body",
            )
        )
        split_boundary_calls = called_names(functions["split_two_taxon_lookup"])
        prohibited_boundary = {
            "graph_two_taxon_lookup",
            "graph_composite_state",
            "graph_composite_recovery",
            "evaluate_split_composite",
        }
        boundary_overlap = sorted(split_boundary_calls & prohibited_boundary)
        rows.append(
            check_row(
                "S03_split_boundary_no_graph_or_shared_composite_calls",
                not boundary_overlap,
                "boundary-specific split lookup",
                ",".join(boundary_overlap) or "none",
                "split boundary lookup performs its own empirical-index query",
            )
        )
        construction_calls = []
        for node in ast.walk(functions["main"]):
            if not isinstance(node, ast.Call) or not isinstance(node.func, ast.Name):
                continue
            if node.func.id != "restriction_from_k_edges":
                continue
            fourth = node.args[3] if len(node.args) >= 4 else None
            construction_calls.append(
                isinstance(fourth, ast.Name) and fourth.id == "observed_k"
            )
        rows.append(
            check_row(
                "S04_observed_k_fed_to_explicit_k_helper",
                any(construction_calls),
                "restriction_from_k_edges(..., observed_k)",
                str(construction_calls),
                "construction-order branch must use the observed ordered-anchor K",
            )
        )
        graph_restriction_calls = called_names(functions["graph_restriction"])
        rows.append(
            check_row(
                "S05_graph_restriction_delegates_to_explicit_k_helper",
                "restriction_from_k_edges" in graph_restriction_calls,
                "delegates",
                str("restriction_from_k_edges" in graph_restriction_calls).lower(),
                "expected restriction and observed-order restriction share only the supplied-K derivation helper",
            )
        )
        recurrence_names = (
            "labelled_set_partitions",
            "rooted_partition_tree_count",
            "unrooted_topology_count_recurrence",
        )
        recurrence_calls = set().union(*(called_names(functions[name]) for name in recurrence_names))
        forbidden_generation = {
            "binary_topologies",
            "all_normalized_topologies",
            "contract_internal_subset",
            "topology_signature",
            "insert_leaf",
            "empirical_split_index",
        }
        recurrence_overlap = sorted(recurrence_calls & forbidden_generation)
        rows.append(
            check_row(
                "S06_recurrence_separate_from_topology_generation",
                not recurrence_overlap,
                "no generator calls",
                ",".join(recurrence_overlap) or "none",
                "set-partition recurrence is executable without graph topology generation",
            )
        )

    failed = [row for row in rows if row["status"] != "PASS"]
    audit = {
        "status": "PASS" if not failed else "FAIL",
        "check_count": len(rows),
        "failed_check_count": len(failed),
        "validation_script": script.as_posix(),
        "checks": rows,
        "scope_note": "AST checks establish prohibited-call separation; scientific independence also requires rowwise output agreement and manual code review.",
    }
    (output / "STATIC_INDEPENDENCE_AUDIT.json").write_text(
        json.dumps(audit, indent=2, sort_keys=True) + "\n", encoding="utf-8"
    )
    with (output / "STATIC_INDEPENDENCE_CHECKS.tsv").open(
        "w", encoding="utf-8", newline=""
    ) as handle:
        writer = csv.DictWriter(
            handle,
            fieldnames=("check_id", "status", "expected", "observed", "detail"),
            delimiter="\t",
            lineterminator="\n",
        )
        writer.writeheader()
        writer.writerows(rows)
    print(json.dumps({"status": audit["status"], "checks": len(rows), "failed": len(failed)}, sort_keys=True))
    return 0 if not failed else 1


if __name__ == "__main__":
    raise SystemExit(main())
```
