## Supplementary Computational Evidence for "A Unique Graph-Theoretic Truth Table for Cross-Gene Branch Identity": VALIDATION_PROTOCOL_v0.2.md

# Computational validation protocol v0.2

Status: **FROZEN BEFORE IMPLEMENTATION**  
Date: 2026-07-23 JST

## 1. Frozen scientific domain

The scientific domain is unchanged:

- Reference tree: `(A,(B,C),(D,(E,F)));`.
- Main retained sets: every subset of `{A,B,C,D,E,F}` of size three through six, 42 sets.
- Main empirical topologies: every normalized labelled unrooted topology, including multifurcations.
- Expected topology counts by retained-taxon count: 1, 4, 26, and 236; these values must also be produced by an executable set-partition recurrence separated from topology generation.
- Expected main cases: 472, comprising 260 binary and 212 multifurcating cases.
- Primitive cells: 4,248.
- Composite axis: 21 prespecified member sets; composite cells: 9,912.
- Total main categorical cells: 14,160.
- Construction-order trials: 1,920; suppression-order trials: 84.
- Certified boundary domain: 15 two-taxon and 6 one-taxon retained sets.

No additional reference tree, empirical dataset, topology class, or manuscript claim is introduced.

## 2. Implementation requirements

### 2.1 Independent split-side composite evaluator

The graph-side composite evaluator continues to consume graph-derived deleted coordinates and original-edge fibers. Its graph-side recovery evaluator consumes graph-derived component splits and the empirical split index.

The split-side evaluator must be a separately named, self-contained implementation. It consumes only split-derived structural coordinates, projected-split groups, the group-to-split index, a composite member set, the empirical split index, and declared split lengths. It must not call the graph-side `composite_state` or `composite_recovery` functions and must not reuse their returned objects. It must derive:

- `active`, `decomposable`, or `unavailable` from split-group coverage and structural/group-partial-intersection rules;
- selected split components directly from the split-group index;
- `recovered_active`, `recovered_decomposable`, or `unrecovered` by actual empirical-index queries;
- `single_edge_lookup`, `selected_component_edge_sum`, or `no_numeric_value`;
- the conditional numerical value.

Shared elements are limited to immutable primitive data, the common empirical input, and canonical serialization/sorting helpers. A separate AST-based check must confirm the absence of calls from the split-side evaluator to the two graph-side composite functions.

Every one of the 9,912 composite rows must compare graph-side and split-side state, component member sets, component splits, recovery state, construction mode, and conditional numerical value. Agreement may be reported only from persisted rowwise comparisons.

### 2.2 Explicit two-taxon empirical recovery

For each two-taxon retained set, construct a normalized empirical graph consisting of the two labelled leaves joined by one empirical edge. Build its empirical split index using the same general split-index constructor used for main empirical trees. Assign the single empirical split the declared deterministic valid length `Decimal("1")`.

The graph-side boundary lookup must derive the path member set from the graph restriction, derive its projected split, query the explicit empirical split index, and return state/recovery/mode/value. The split-side boundary lookup must independently derive the corresponding group and projected split from the split restriction and query the same explicit empirical input without calling the graph-side boundary or composite evaluators.

Persist the empirical topology signature, empirical edge count, lookup split, declared length rule, graph and split reference states, graph and split recovery states, graph and split modes, graph and split numerical values, and row pass status. Discrepancies must be recorded for any disagreement or failed lookup.

### 2.3 Construction-order fiber derivation from observed K

Create a helper that accepts an explicitly supplied `K` edge set and derives deleted coordinates, degree-two adjacency, original-edge fibers, coordinate-to-fiber mapping, and primitive reference states from that exact edge set. The normal all-pairs graph restriction must compute its expected `K` and delegate to this helper.

For each of the 1,920 construction orders, compute `observed_k` from that order, pass `observed_k` to the explicit-`K` helper, and directly compare both:

1. `observed_k` versus expected all-pairs `K`; and
2. fibers derived from `observed_k` versus expected fibers.

The persisted order table must expose both comparisons separately in addition to the overall pass flag. A static check must confirm that the construction-order branch supplies the variable `observed_k` to the explicit-`K` helper.

### 2.4 Independent set-partition recurrence

Implement an executable recurrence that is separated from binary insertion, edge contraction, topology signatures, and `all_normalized_topologies`.

Let `R(1)=1`. For a labelled set `S` with `|S|>=2`, define

`R(S) = sum over set partitions pi of S with |pi|>=2 of product over B in pi of R(B)`.

Rooting an unrooted normalized tree at the neighbour of a distinguished removed leaf gives `U(n)=R(n-1)`. The recurrence must therefore produce `U(3..6)=1,4,26,236`. Generated topology counts are observed values and recurrence counts are independent expected values. Persist the reconciliation in a dedicated machine-readable table. Output-only QA must independently recompute the recurrence without importing the validation implementation.

## 3. Pass/fail and discrepancy policy

The validation passes only if all of the following hold:

- all frozen domain counts reconcile;
- all 4,248 primitive comparisons pass;
- all 9,912 strengthened composite comparisons pass;
- all 1,920 construction-order rows pass both the `K` and observed-`K-derived fiber` comparisons;
- all 84 suppression-order rows pass;
- all 15 explicit two-leaf empirical lookups agree on state, recovery, mode, and value;
- all 6 one-taxon boundary cases retain their certified categorical behavior;
- the independent recurrence agrees with generated counts for sizes three through six;
- every required state and class remains covered;
- `DISCREPANCIES.tsv` is present with a header and contains zero data rows only if the rowwise derivations produce no discrepancy;
- the static code-path audit passes;
- two clean runs are byte-identical for every canonical scientific output;
- output-only QA passes all declared checks;

Any failure forbids a success claim or manuscript-ready zero-discrepancy claim. Failed rows, logs, and discrepancies must be retained truthfully.

## 4. Claim boundary

The strengthened computation checks implementation fidelity on the prespecified finite six-taxon domain. It does not prove the theorem, certify production SplitAligner, expand the certified boundary scope, or make the computation a premise of the theorem. The computation remains independent of the mathematical proofs and supports only the stated finite-domain implementation-fidelity claim.
